# The cavity method for community ecology

**DOI:** 10.1101/147728

**Authors:** Matthieu Barbier, Jean-François Arnoldi

## Abstract

This article is addressed to researchers and students in theoretical ecology, as an introduction to “disordered systems” approaches from statistical physics, and how they can help understand large ecological communities. We discuss the relevance of these approaches, and how they fit within the broader landscape of models in community ecology. We focus on a remarkably simple technique, the cavity method, which allows to derive the equilibrium properties of Lotka-Volterra systems. We present its predictions, the new intuitions it suggests, and its technical underpinnings. We also discuss a number of new results concerning possible extensions, including different functional responses and community structures.

## 1 Context

### 1.1 Disordered systems

Many foundational insights of ecological theory were derived from the careful study of simplified patterns of interaction between a few distinct entities (individuals or species). Examples include trophic chains [20], competition and exclusion over a few resources [21, 39], mutualism [13] and parasite-host interactions [25]. These archetypal structures can be used as modules [15, 5] to understand more complex systems, when this complexity does not interfere with their order and structure. For instance, our intuitions on multitrophic communities combine simple trophic mechanisms to understand inter-guild relations, and simple competitive mechanisms for intra-guild dynamics. Many theories also distinguish between ecosystems shaped by top-down control from predators, and those shaped by bottom-up control from resources. In all cases, these intuitions can only work if these interaction structures remain distinct even in a complex community.

In parallel, neutral models [16] have shown that many quantitative predictions can be made with minimal assumptions of structure, or even of distinctiveness of the entities. In most such works, the only accounting for an ecosystem’s diversity is its number of otherwise identical species. These theories belong to the *mean-field* class, meaning that intrinsic differences between entities are erased statistically, owing to large numbers and some form of stochasticity. More precisely, such differences lose their importance in the collective dynamics – and therefore also in individual species’ dynamics, provided that those are mostly shaped by collective factors (e.g. aggregate competition from many other species).

*Disordered systems* stand between these extremes, in that they exhibit heterogeneity without order: they represent situations where one cannot neglect the existence of differences between species, but no clear-cut structural patterns can be identified. As in neutral theories, it is either impossible or unnecessary to assign clear “roles” to individual species. But instead of being modelled as identical, all species appear to be completely heterogeneous in their biological features and interactions, without regular motifs. What we can know is the range and frequency of these traits in the community as a whole, and we investigate how much can be predicted about an ecosystem from that limited knowledge. While the notion of disordered system originates in physics^1^, perhaps the simplest analogy is found in models of neural networks [27]. Each neuron has a distinct set of interactions, but it is difficult to identify the role of any given neuron, or group them into functional classes; mainly, we can access summary statistics of “how different” neurons and their interactions are in a given network.

This approach shares a conceptual starting point with Random Matrix Theory, which has long become a staple of theoretical ecology [24, 1]. In both cases, one considers a “simple sort of complexity”: in large disordered communities, most details cease to matter and a few general mechanisms emerge, allowing us to unify many superficially different systems and models. However, Random Matrix Theory has developed with a specific focus on stability, whereas disordered systems techniques give access to properties – functioning, coexistence, abundance distributions, trait selection – which are usually predicted using neutral or structured models. In fact, many ecological models already fall into this category, especially numerical models that contain many species with uncertain attributes (lacking empirical or mechanistic constraints) such as large random Lotka-Volterra communities. What has perhaps not been sufficiently realized is the availability of mathematical techniques for making broader and more robust statements about such systems, thus helping guide, validate or even bypass heavy simulation work. It is only recently that these techniques and ideas have been brought to bear on ecological problems [12, 18, 6, 37, 38].

Of particular interest is the application of this framework to *ecosystem assembly,* i.e. how dynamics and interactions shape the realized community from a pool of potential invaders. Indeed, it is easier to imagine that the pool itself is disordered: it may consist of species immigrating from a number of different environments, with independent evolutionary histories. By contrast, the assembled state that emerges from that pool will be less random. By following the process of assembly from that pool, it is possible to see which properties are selected by the ecological dynamics, i.e. how traits in the assembled community differ from those in the pool. Multiple works have thus explored what shapes species coexistence and invadability, in the context of assembly from a fixed regional pool [6], from immigration and stochastic extinction in an island biogeography perspective [18], or from two consumer communities “coalescing” – starting to compete on the same resources [37]. These methods have proven apt at generalizing to many species what was previously understood for modules of a few heterogeneous species, e.g. the conditions of competitive coexistence [38].

### 1.2 Models of ecological dynamics

#### 1.2.1 The disordered systems framework

Practically speaking, the notion of disordered systems, and the fact that it allows for a simple and generic analytical approach, comes with significant methodological potential: it connects a class of commonly used models in ecology, synthesizing their insights and explaining why and when they may give similar or different predictions. We illustrate this in another work [3] with a “model meta-analysis” demonstrating that a variety of community structures, including mutualistic networks and resource competition, can still fall within the realm of disordered systems and create assembly patterns that we can predict with this framework.

This also means that if there were reasons to outright reject our results as a valid representation of ecological reality, then an entire class of models would become disqualified at once, as well as many qualitative arguments based on certain notions of complexity. That would be a useful result for ecological theory, limiting the field of investigation in models and simulations.

More concretely, to help situate this framework among preexisting work in theoretical ecology, we show in Fig. 1 a bird’s eye view of the main types of dynamical models we identified in the field. They roughly fall between three extremes. First, fully independent populations, where we use physiological and behavioral traits to predict existence and functioning within a given environment. Second, fully dependent (i.e. very strongly interacting) populations where, for instance, the coexistence of competitors requires precise tradeoffs between their traits. Third, fully stochastic models, where the characteristics of species and their interactions are all blurred and dominated by noise coming from internal (demographic) or external (environmental) sources. On this graph, our position is represented by the orange dot, a regime of collective organization where interactions are individually weak but collectively strong.

**Figure 1:**
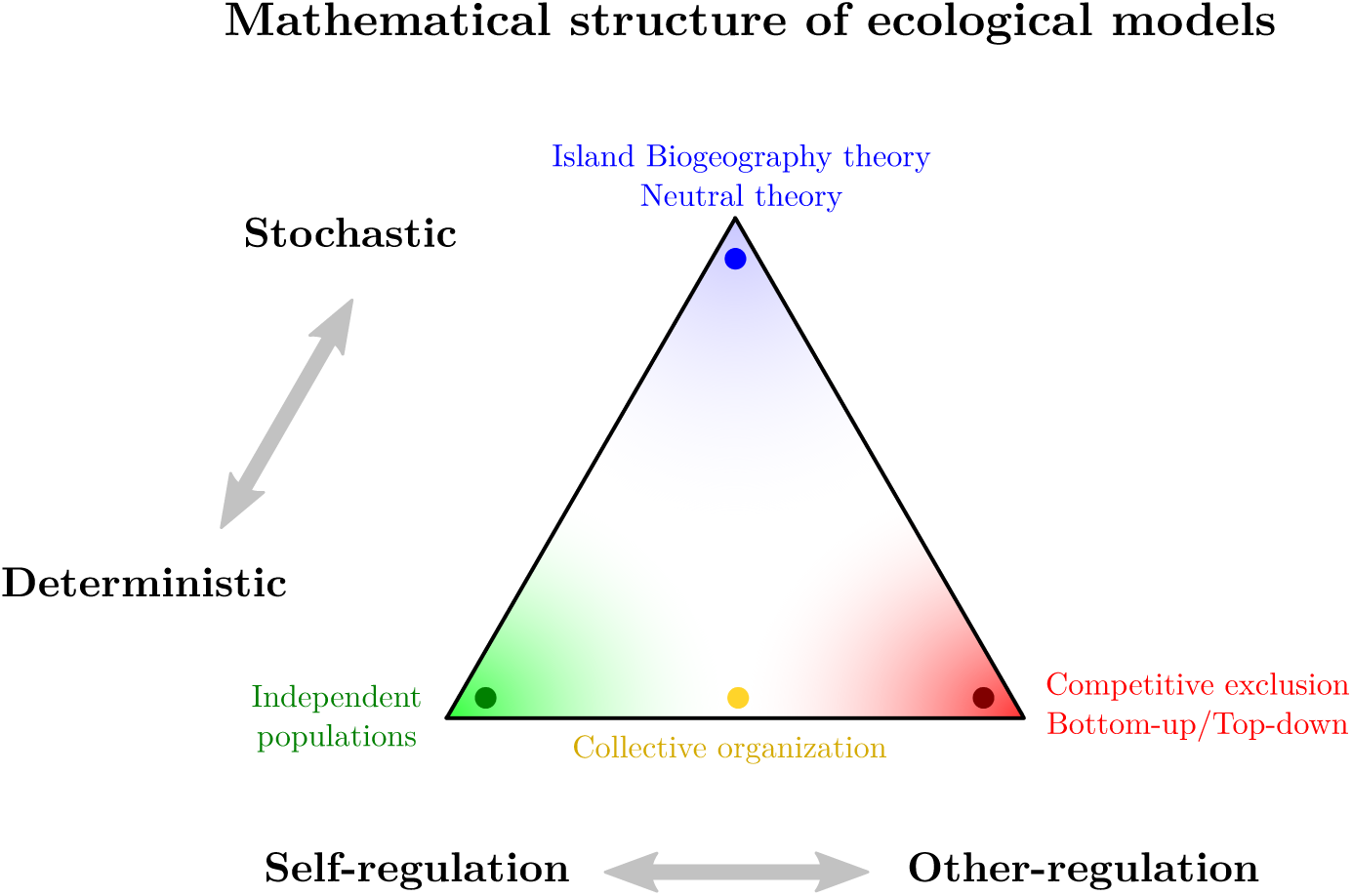
An abstract representation of the main classes of dynamical models in theoretical ecology, and the regime of collective organization where the notions and methods developed here are meant to apply.

The two main axes of opposition on this graph are the following. First, deterministic versus stochastic: in deterministic approaches such as ours, we generally expect that real ecosystem properties can be understood from the equilibrium or near-equilibrium properties of a given dynamical model, controlled by some combination of intrinsic species fitness and by their interactions. The opposite pole includes theories that are dominated by stochastic and transient dynamics, where “noise” – in the demographics, the environment, or the immigration process – prevents the system from ever being close to some deterministic assembled equilibrium^2^. Those include neutral theories [16] and island biogeography and its variants [22, 11, 18]. These two viewpoints are complementary; only careful confrontation to empirical data can decide whether an ecosystem’s dynamics are governed by transient, far-from equilibrium dynamics such as neutral turnover, or by the sort of near-equilibrium features described here.

The second axis of opposition confronts models whose dynamics are dominated by species characteristics, and those that are determined by interactions. Independent populations illustrate the former, competitive exclusion the latter. In these two extremes, results are often very sensitive to model assumptions: for instance, the colonization-competition tradeoff model ensures global coexistence by requiring a very precise ordering of competitive and dispersive ability. It may be surprising to see that the middle-ground – collective organization, where interactions are individually weak but collectively strong (because distributed among many species) – gives rise to very generic and predictable patterns.

This is the class of systems that we can describe with this framework. Note however that it could, fairly easily, be extended to include noise; by contrast, systems where species are strongly influenced by other individual species tend to have very dramatic dynamics (e.g. predator-prey cycles) which typically preclude the existence of any non-trivial equilibrium. Such dynamics are not entirely beyond the reach of disordered system approaches, but will almost certainly require more involved techniques than the simple one presented here.

#### 1.2.2 Example: History dependence

Importantly, each class of models makes different predictions for the mechanisms of various ecological phenomena. We can illustrate it in the case of history-dependence (e.g. founder effects). In a deterministic equilibrium viewpoint, the long-term state of a system is dependent on its history only if its dynamics allow for multiple attractors. Of course, even with a single global attractor (e.g. a single final, stable and uninvadable community), if the time to reach it is very long, the system may retain some memory of its previous states at visible timescales. But this memory should become fainter as time passes, and any shock should only reset the dynamics to some earlier stage along a similar trajectory. This perspective is compatible with the intuitive notion of maturity^3^ as found e.g. in Margalef [23].

By contrast, in a stochastic perspective, any composition is to some degree an accident, though it may hold for a long time if there are positive feedbacks (e.g. self-reinforcing density dependence) that transform an early advantage into a long-lasting one. Nevertheless, the composition may well undergo a dramatic shift, due to a shock or even out of steady noise, without it signalling any change in intrinsic species fitness. This is closer to the usual intuition behind the founder effect.

## 2 General framework

### 2.1 The assembly process

Here is the central conceit of this theory of assembly: we define a pool of species that can invade the community, and whose dynamical traits, such as growth rates and interactions, are known only statistically. We then allow sequences of invasions until the community reaches an uninvadable state, and investigate the properties of this “assembled” state – among others, species diversity, abundance, and how traits selected by the dynamics differ from the pool baseline.

This can represent a variety of physical settings: the pool could be a entire region and the community a local patch [31], they could be a mainland and an island connected by migration fluxes [22, 11, 18], or even oceanic water and the microbiome of an individual sponge [36]. More generally, this suggests a stable reservoir of biodiversity with some known aggregate properties, and an environment that is being populated from that reservoir, even if the reservoir is not larger (e.g. a forest serving as a biodiversity shelter for surrounding plains), provided that it can serve as a stable supply of potential invaders for the timescales considered.

But this can also represent, more abstractly, a “separation of concerns” between, for example, physiology and ecology. Properties of the pool reflect the traits of species “in potential”; by contrast, the assembled community retains only the subset of traits that are dynamically selected to survive together. Thus, in an abstract model, the pool could represent the range of species that are allowed to exist by physiological constraints, while assembly would single out combinations of such species that can coexist in the same ecosystem. Even in the spatial interpretation suggested before, there is such a separation of concerns: species-species interactions are properties of the pool, but species-environment interactions (e.g. carrying capacities) could be specific to a location, leading to different assembled states in different locales.

This means that the size of the pool, *S*, admits very different interpretations depending on the ecological scenario represented: *S* could be regional diversity or mainland diversity, if one is interested in invasion through immigration, but it could also represent the mutant strains that try to invade a microbial community over a given timescale (the pool being disordered would then reflect an assumption of random mutations). Carefully teasing apart these various interpretations and the hypotheses they hide is one of our purposes in this article.

### 2.2 The assembled state

#### 2.2.1 Equilibrium relations

Within this general setting, we decide to look only at the final assembled state, i.e. a feasible, stable and uninvadable community. As discussed in Sec. 1.2.1, this stands in stark contrast with theories that are dominated by stochastic and transient dynamics.

The mathematical techniques of disordered systems can be applied to many different dynamics. For convenience, we investigate the very commonly used Lotka-Volterra dynamics^4^:

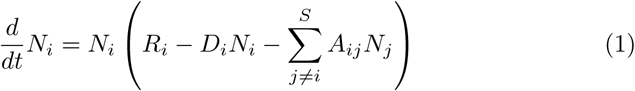

Here *A_ij_* is the interaction between species *i* and *j* (i.e. the rate at which a unit of species *i* affects a unit of species *j* dynamically). *R_i_* is the growth rate of species *i*, that combines growth on external resources and mortality (and could thus be negative). *D_i_* is intra-species competition, hereafter *self-interaction.* We can define

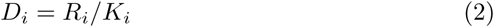

where *K_i_* is properly the carrying capacity of species *i* in the case of consumers with *R_i_* > 0. If *R_i_* < 0, then *K_i_* is negative and cannot be interpreted as a carrying capacity, but remains an important mathematical quantity (a measure of density-dependent mortality).

We will concern ourselves only with the equilibrium solutions, meaning *dN_i_/dt* = 0 and

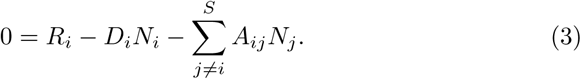

Only the species for which a positive solution *N_i_ >* 0 exists will then survive together in an assembled state. This state is obtained after long immigration sequence, during which all species that could invade did. As we explain below in Sec. 3.3, in a certain region of parameters, the precise sequence of invasions is irrelevant to the final state, which is a global attractor of the dynamics.

#### 2.2.2 Control parameters

**Important: To simplify the exposition, we assume starting now that all the self-interaction terms are equal** *D_i_* = 〈*D*〉 = 1, **so that** *K_i_* = *R_i_.* **This assumption is relaxed and the consequences of heterogeneous** *D_i_* **are discussed starting in Sec. 4.**

We will proceed to show that, to predict many properties of the assembled state, it is sufficient to know four simple statistical features of the pool of potential invaders. First, we need a measure of intrinsic fitness differences between species; for this we take *ζ* the standard deviation of the growth rates *R_i_.* Second, we need to characterize interactions at a collective level. We thus ask how much a species is affected by the *sum* of all its interactions: how adverse this total interaction is on average, represented by a parameter *μ* (*μ >* 0 means that biomass is on average lost through interactions, *μ <* 0 means it is gained), and how much that varies from species to species, represented by a parameter *σ.* Hence, for any species, the total effect of interactions can be understood to fall roughly in the range *μ* ± *σ.* Finally, to understand feedbacks between species, we need to know how similar the effect of species *i* and *j* is to the reciprocal effect of *j* on *i*, and we encapsulate this in a parameter *γ.*

Formally, these parameters are defined as follows:

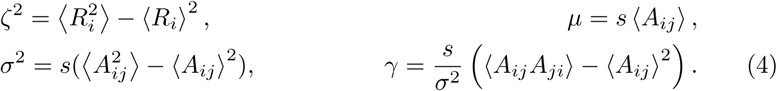

where *s* = *Sc* is the average number of interaction partners for each species (*c* being the connectivity of the interaction network). This factor *s* means that *μ* and *σ*^2^ indeed represent the *total* interaction strength and variance faced by one species from all its partners. Given our definition of *γ*, it takes values in the range [−1, 1]: *γ* = 1 means that two partners affect each other identically, while *γ* = −1 means that reciprocal effects are maximally different, see Sec. 4.2.2. Examples from competitive and trophic communities are given in the next section.

These parameters suggest how disordered systems differ from both mean-field and structured systems. The various species traits retain a non-negligible heterogeneity, encapsulated by *ζ* and *σ.* Yet, little more information is needed than those simple metrics, which give a basic estimate of “how different” species can be. The precise interpretation of these abstract parameters will become clearer, first from seeing what role they play in model results, and later in more in-depth discussion.

All higher-order correlations, that capture more precise structural information about the pool, are neglected. If more order is needed (e.g. for trophic systems, given that trophic roles are an essential feature that is not represented here), one could consistently push the formalism further, incorporating more and more structural properties in the form of correlations^5^.

#### 2.2.3 The special case of competitive communities

Note that in competitive communities, where species have carrying capacities *K_i_ >* 0 which can follow a wide distribution (e.g. lognormal), it is often difficult to generate communities where many species coexist if interactions *A_ij_* are drawn arbitrarily. It has been found that a particualr scaling of interactions does allow for species coexistence: they must scale with the ratios of carrying capacities, more precisely

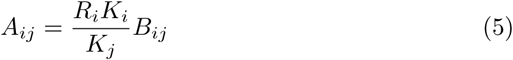

where *B_ij_* has a narrower distribution (e.g. normal) [17, 26]. In this case, we can divide the equilibrium condition (3) by *R_i_* to get

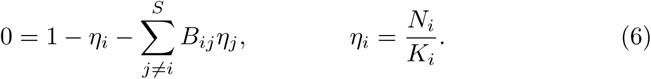

In other words, all the calculations and results presented here apply to this particular competitive setting by replacing the *N_i_* with the effective variables *η_i_*, the real interactions *A_ij_* by their “core” *B_ij_*, and setting *R_i_* = *D_i_* = 1.

Our framework in fact gives a very clear mathematical explanation for how the scaling s(5) allows for coexistence: imagine instead that we had started from the Lotka-Volterra equations rewritten as

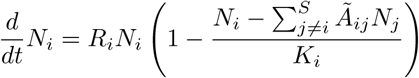

with some randomly drawn *Ã_ij_*, then the equilibrium condition would be

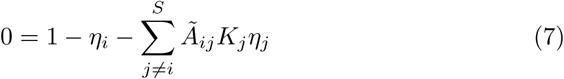

for the effective variables *η_i_* = *N_i_/K_i_*. The effective interactions *B_ij_* = *Ã_ij_K_j_* could be very widely distributed, since *K_j_* themselves are drawn from some wide (e.g. lognormal) distribution. This would correspond to a large value of interaction heterogeneity *σ*, which we show below has a very negative effect on coexistence. We give precise conditions on the statistics of *B_ij_* which allow to generate communities of given diversity.

## 3 What can we predict?

While the following sections discuss the methods that we will use to solve the equations above, this one gives an overview of some of the main system properties that can be computed in this framework. These results fall roughly in four categories of important ecological properties: functioning, diversity, stability, and features selected by the assembly process.

We present these results in the abstract parameter space defined in (4). The interpretation of the parameters comes in Sec. 4. Importantly, the results that we show are exact for the region of parameters where there exists a single, globally stable, uninvadable final assembled state. However, most results hold approximately past the boundary of this region, and we later discuss in Sec. 4.1 when systems are likely to be found in this region. The complex region that lays beyond is still being investigated.

### 3.1 Functioning

For functioning, we can for instance compute the total biomass

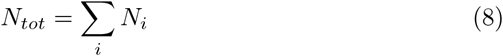

and total productivity or growth from external resources

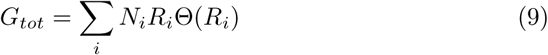

with the Heaviside function Θ(*x*) = 1 if *x >* 0, Θ(*x*) = 0 otherwise. In fact, since *G_tot_* scales with the total biomass, it is more practical to look at the productivity ratio *G_tot_*/*N_tot_*, which increases when faster-growing species are favored.

Note that throughout, we make no difference between biomass and abundance, as we are not considering individual mass as a particular species trait. Our choice is equivalent to assuming that biomass has no systematic correlation with either interactions or carrying capacities in the pool of potential invaders, so that converting total abundance to total biomass only requires multiplying by the average individual’s mass in the community. However the framework could easily be extended to account for individual mass as a trait controlling other dynamical properties (e.g. via metabolic scaling).

In addition, let us recall here that, as suggested in Sec. 2.2.3, the calculations made here can be extended to competitive communities with wide distributions of carrying capacity *K_i_*, but all our predictions for *N_i_* will instead be about the rescaled abundances *N_i_/K_i_*. Hence, for instance, predictions of 〈*N*〉 *>* 1 will signal overyielding. Since these discussion points are specific to competitive communities and we strive to be as general as possible, we will leave the analysis of all these consequences to future work.

### 3.2 Diversity and coexistence

For diversity we will consider two main metrics. The first is the number of species surviving in the assembled state *S\** = *Sø*, see Fig. 2. As one would expect, coexistence decreases with increasing *σ* and *ζ*, since more variance in growth rates and interactions entails that some species are more likely to be overall stronger competitors than some others. Interestingly, as we discuss below, decreasing *γ* tends to increase coexistence. The second is Simpson’s index, which gives a measure of biomass concentration,

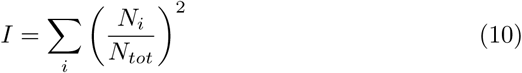

**Figure 2:**
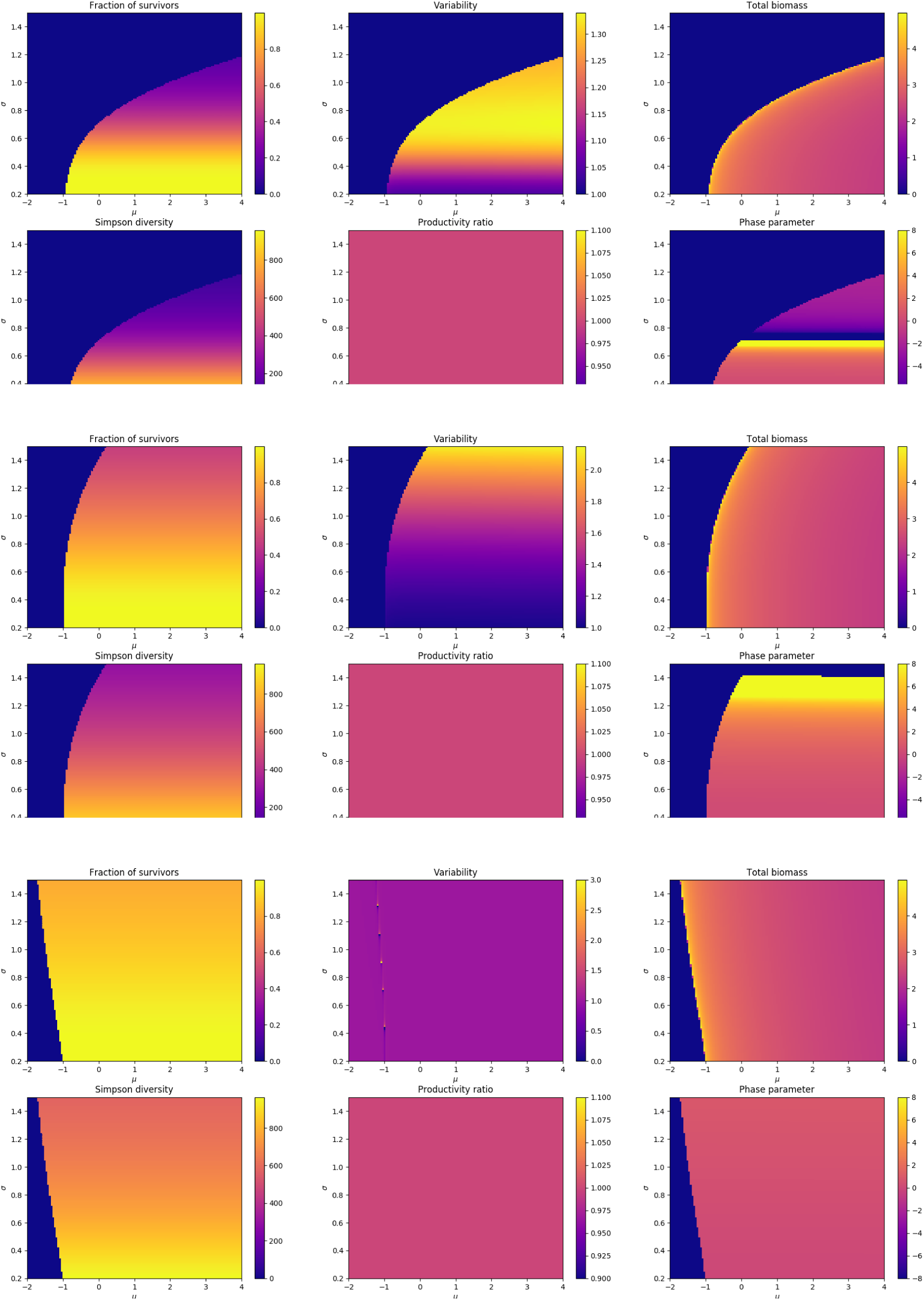
Coexistence, abundance and stability properties in the space (*μ, σ*) for *γ* = 1, 0, −1 (top to bottom) and *ζ* = 0. Since all species have the same growth rate, the productivity/biomass ratio is always constant here. The uniform area in the left of each graph signals the parameter region where some interactions are mutualistic enough that abundances diverge, see Fig. 4 below. As is apparent here, when *ζ* = 0, the average coupling strength *μ* only affects the total biomass *N_tot_,* while all other panels show purely vertical gradients, i.e. they only depend on *σ* (and *γ*). Results for *ζ* = 0.3 are shown below in Fig 5.

Its inverse, Simpson diversity *D*, tells us how effectively diverse the community is, in terms of the biomass being equitably distributed among all species. The index is related to the second moment of the abundance distribution. In fact, the full distribution can be computed within this framework, with some caveats discussed in Sec. 5.

### 3.3 Stability

We discuss three aspects of stability: the existence of multiple equilibria, variability around an equilibrium under stochastic perturbations, and the consequences of permanent community shifts – be they due to environmental changes or species extinction. All three can be shown to be related via simple expressions to pool parameters and to the fraction of surviving species *ϕ*.

First, we can find an expression for *invariability* as defined in [2], i.e. the inverse of the response of the system to demographic noise.

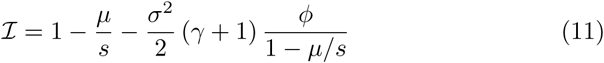

It can formally be related to the response to other types of noise, and is also the basis for deriving a prediction for Taylor’s law.

Second, we can derive the number of secondary extinctions resulting from the extinction of one species in the system. In a disordered system, secondary extinctions are more or less numerous depending on parameters, but never make up a significant fraction of the community, due to species abundances being controlled by the community as a whole rather than by individual partners.

The final important result is *χ* the order parameter for multistability. Let us imagine that we perturb all the carrying capacities

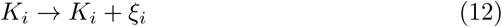

with a perturbation that is on average 〈*ξ_i_*〉 = 0, representing a small change in the environment that leads to different species being favored, without making the environment overall better or worse for the community. Then we can ask how much the abundances of the species are changed in response,

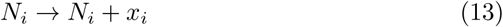

and this change (with zero mean) is characterized by its variance, which we compute in Sec. 5.6 below:

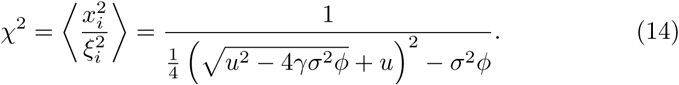

where *u* = 1 − *μ*/*s*. This expression reflects the deep relationship between coexistence *ϕ* and sensitivity to the environment *χ.*

The divergence of *χ*^2^ when the denominator of (14) vanishes, i.e. when

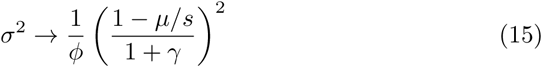

signals the boundary of the region where multiple equilibria exist: precisely at the boundary, an infinitesimal change in intrinsic fitness can lead to different species being present in the final assembled state. Of course, while the quantity formally diverges, the community’s response is not truly infinitely steep – it only signals that the approximations we use to compute the single globally attractive equilibrium break down. Indeed, as one gets close to the divergence, *χ* stops predicting the simulated response to a press perturbation, which plateaus as shown in Fig. 3.

**Figure 3:**
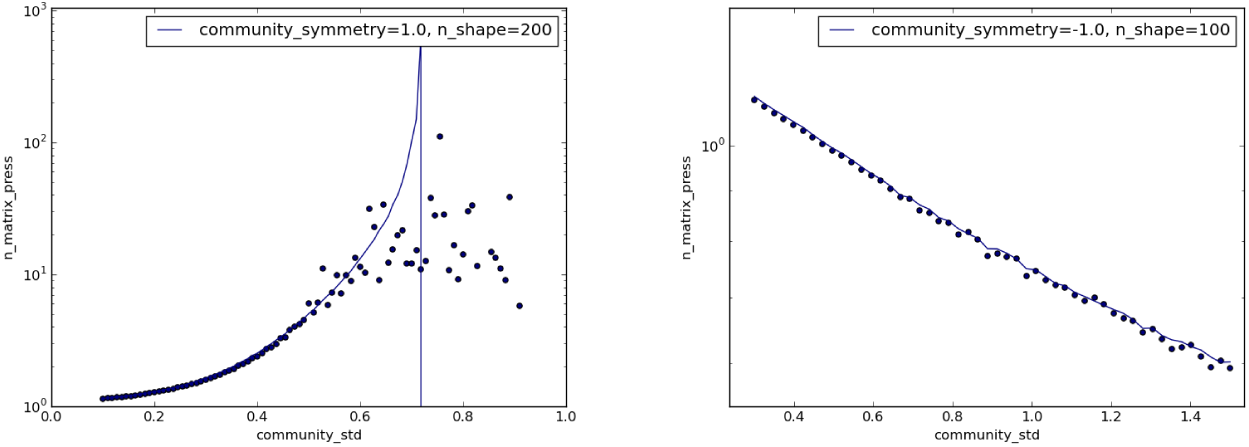
Average effect of a random perturbation of carrying capacities, calculated as *χ* (solid line) or directly from the assembled community in simulations (dots), depending on coupling heterogeneity *σ.* Left: *γ* = 1, with divergence of *χ* at finite *σ,* revealing the loss of stability of the single global attractor and the start of the multistable phase. Right: *γ* = −1, no divergence (no multistable phase).

Hence, it is possible to construct the *phase diagram* shown in Fig. 4, which indicates where in parameter space one will find a single, multiple or no equilibria. As we discuss more below, lowering *γ* has a stabilizing effect, as it extends the single equilibrium phase – in fact, it is apparent from (15) above that, in the limit *γ* = −1, the threshold to the multiple equilibria phase becomes infinite, and there can only be one attractor which is globally stable.

**Figure 4:**
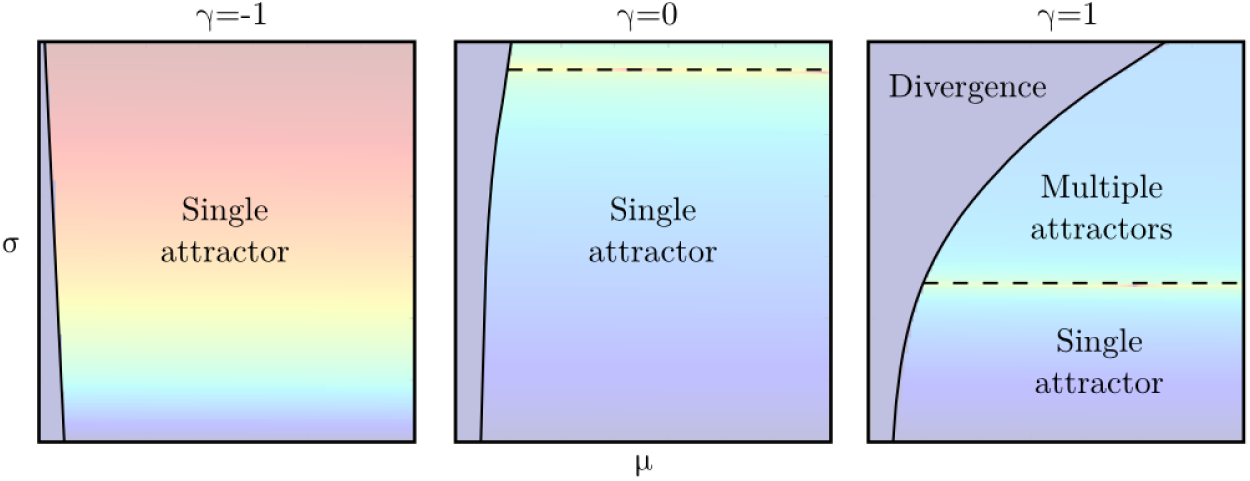
Phase diagram in the (*μ*, *σ*) plane for various *γ* and *ζ* = 0, based on phase parameter *χ*^2^ from Sec. 3.3. There are three regions depending on parameters. In the first region, there is a single global attractor, meaning that – after a long enough sequence of invasions and extinctions – a given community will reach a single uninvadable state, no matter what sequence it went through. In the second region however (up of the dashed line, signalling a divergence of *χ*^2^), there are multiple such attractors, and the final assembled state depends on the history of the system. As *γ* → −1, the transition to the second region disappears. Finally, the third region (left of the solid line, signaling a Rdivergence of 〈*N*〉) has no attractor, as some species have mutualistic interactions that are stronger than any negative feedback and their abundances grow to infinity. This signals a breakdown of the Lotka-Volterra approximation for more realistic dynamics.

### 3.4 Selected features

Previous work [6] has shown that it is possible to use this framework to infer not only the distributions of abundances, growth rates and interactions in the assembled state, but also correlations between those, including the frequency of various small network motifs [15]. How these distributions differ from the pool baseline tells us how non-random properties have been selected from a random pool. This gives us worthwhile information on which factors are likely to enable a species to survive into the assembled state, or conversely, how ecosystems may adapt to ensure coexistence. We demonstrate this idea in this article with the selection of carrying capacities or growth rates, but see the Appendices in [6] for details on the selection of interaction patterns.

### 3.5 Discussion

Most striking is the role of *γ* the reciprocity of interactions: as *γ* becomes negative, sensitivity to perturbations decreases, the single attractor region expands and coexistence increases. This suggests that asymmetrical interactions – including commensalism, amensalism and trophic interactions – lead to easier assembly than symmetrical ones, echoing the conclusions of other authors [1] who noted “remarkable differences between predator-prey interactions, which increase stability, and mutualistic and competitive, which are destabilizing”. Yet, we can also show that *γ <* 0 may very well be achieved by competitive interactions. It is in fact possible to construct trophic and competitive communities so that the latter have more negative *γ* than the former. Hence, this statement should not be taken as generally valid, and the meaning of *γ* should be discussed much more carefully, as we do in Sec. 4.2.2.

In the single attractor region, many different ecological scenarios become indistinguishable as long as one is only interested in the eventual assembled state. Whether invaders from the pool come from migrations or mutations, frequent or rare, correlated or not with the species dominating the community at any given time, interspersed with extinctions and recoveries or not – these distinctions become irrelevant in the long run, as any dynamical trajectory that does not involve irreversible changes in the pool of potential invaders (e.g. a permanent extinction) will end at the same point.

If we look at a pool that lies on the boundary between the two regions, multiple attractors first emerge as small deviations from the single attractor that existed below that boundary - most species remain, and only a few rare ones distinguish the multiple attractors by their absence or presence. Therefore, even though the results presented in this work focus on the single attractor region, where our analytical results hold exactly, we will demonstrate multiple times that our framework still provides good approximations some distance into the multiple attractor region. When our framework ceases to be predictive altogether, it is likely that the various attractors are different enough that perturbations may engender significant regime shifts.

If *ζ* ≠ 0, the model predictions seen on Fig. 5 are still quite similar to those shown in Fig. 2 for *ζ* = 0, except for the fact that average coupling *μ* plays a much more important role: instead of controlling only the average biomass 〈*N*〉, it affects all the metrics, favoring the existence of a global attractor while reducing coexistence.

**Figure 5:**
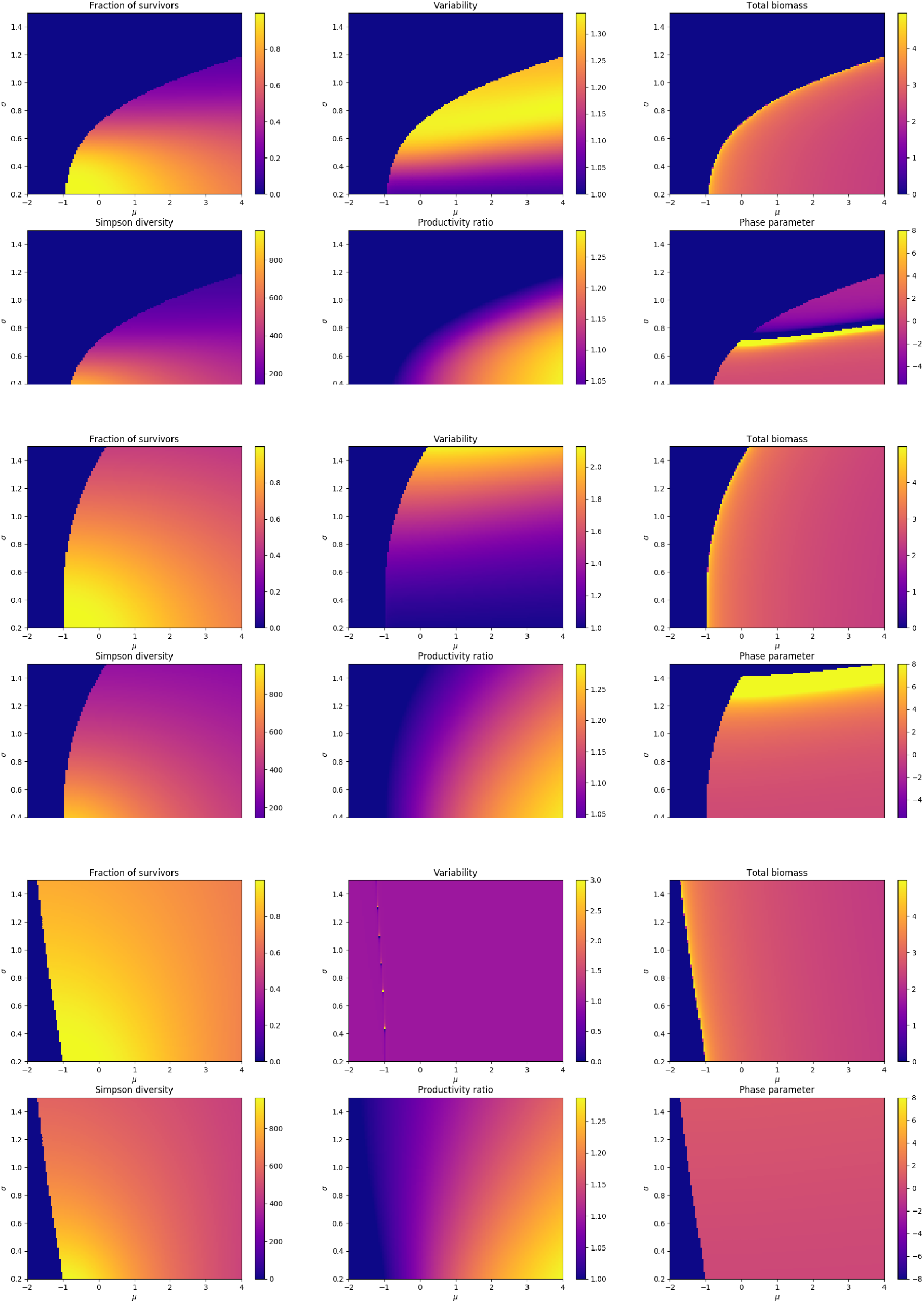
Coexistence, abundance and stability properties in the space (*μ*, *σ*) for *γ* = 1, 0, −1 (top to bottom) and *ζ* = 0.3. See Fig. 2 for results at *ζ* = 0.

## 4 Mathematical setting

This section is meant to be an qualitative explanation of the mathematical concepts that are crucial for a better understanding of the methods and results shown in this paper. Sec. 5 below gives a detailed explanation of the derivation, of interest to the more mathematically-inclined reader.

We recall that the central equation we investigate is (3) which defines the equilibrium for Lotka-Volterra dynamics:

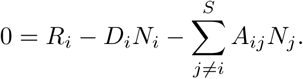

Before solving this equation, we show that it is possible to better isolate the role of the various parameters, thanks to two simple tricks of rescaling. First, it is easily seen in (1) that *R_i_*, *D_i_* and *A_ij_* could all be multiplied by the same number, without changing the dynamics (only making them globally faster or slower). Hence we can arbitrarily choose the time scale so that, for instance, growth rates have unit mean:

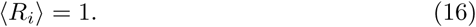

Now, we can look more specifically at equilibria, i.e. equation (3). Clearly, we could multiply all *D_i_* and *A_ij_* by some number, and divide all *N_i_* and *N_j_* by the same number, without changing the equilibrium: the average intensity of interactions, inter- and intra-species, simply changes the average abundance at equilibrium. Hence we can also arbitrarily take

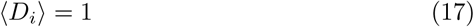

so that now, interactions *A_ij_* are measured in units of the average self-interaction, and abundances are measured in units of “carrying capacity” – indeed, if it were alone with positive growth rate, the average equilibrium abundance of the species would be around 1 (or equal to 1 if *D_i_* = 1). Neither of these choices subtract anything from the generality of the equations. An important consequence is that any change of the absolute value of growth rates or interactions, for instance via nutrient enrichment, will affect only the total amount of biomass and no other property of the assembled state.

The second trick is using the fact that interactions can always be rewritten as

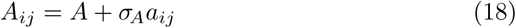

with two parameters: the average interaction *A* = 〈*A_ij_*〉 and the variance of interactions 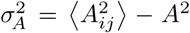, so that the rescaled interactions *a_ij_* have zero mean and unit variance

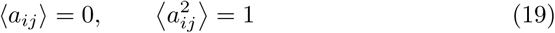

For now, we make no further assumption on what the distribution of the *A_ij_* and *a_ij_* may be, beyond the fact that it has a well-defined mean and variance. We simply suggest that is important to know both how intense interactions are on average, i.e. *A*, and how much they differ between pairs of species, *σ_A_*.

Hence, we transform (3) into the following equation

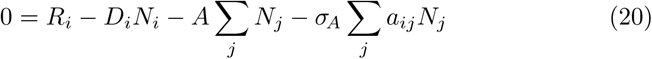

and using

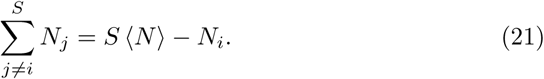

we finally get

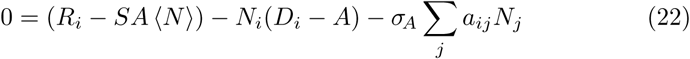

The first term is an effective growth rate including the overall biomass loss due to competition (*A >* 0) or gain due to mutualism (*A <* 0). The second term contains the effective intra-species competition, and the third the relative gain or loss of fitness of species *i* due to the *heterogeneity* of inter-species interactions. Multiple insights already arise from this simple reformulation, without even solving the equation, and we discuss them now.

### 4.1 Scaling of interactions: when are large disordered communities not neutral?

#### 4.1.1 Inter-species versus intra-species effects

It is well-known that intra-species competition plays a large role in stabilizing the dynamics: interactions with other species become important only if they are not negligible compared to this self-interaction. Strongly interacting communities can exhibit complex dynamics with multiple equilibria, limit cycles or other attractors. If interactions with others are negligible, each species simply follows its own logistic equation, and the system is sure to evolve toward a globally attractive equilibrium.

The equation (22) above shows that – beyond a simple translation of every species’ growth rate – the effective importance of interactions compared to self-interaction is controlled by the ratio

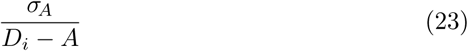

This gives a clear intuition of why species heterogeneity, as encapsulated by *σ_A_*, can be important in the dynamics. If species are too similar, *σ_A_* ***→*** 0, then the abundances *N_i_* decouple from each other: species are only coupled collectively through the average value 〈*N*〉 in the effective growth rate, except in the very special case of the Hubbell point *D_i_* = *A* where all inter- and intra-specific interactions are equal [16, 18]. Away from that singular point, we see that what truly destabilizes the system is not only how *intense* interactions are, i.e. *A* at the denominator, but also how *diverse* they are, i.e. *σ_A_* at the numerator.

#### 4.1.2 Comparing communities of different sizes and connectivities

A second central insight comes from asking: what does it mean to say that two pools with different sizes *S* have the *same amount* of heterogeneity? Let us assume that each species is connected to *s_i_* other species in the interaction matrix, on average

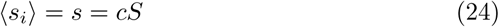

with *c* ∊ [0, 1] the connectivity of the network. If *s* is the same in the two pools^6^, then species from either pool have the same average number of interactions. In that case, it makes sense to directly compare the properties of these interactions, i.e. the values of *A* and *σ_A_*, to decide which pool has stronger or more diverse interactions.

However, let us assume that *c* is the same for both pools, e.g. *c* = 1. In that case, the dynamical influence on one species from all the others will be proportional to the quantity

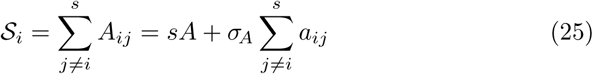

(this gives a general trend; of course, the precise dynamical effect will depend on species abundances). This quantity is on average

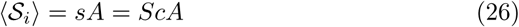

which increases with the size of the community. This term is seen in (22) to intervene in the effective growth rate. As for how much 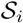 varies between species, that comes from the sum

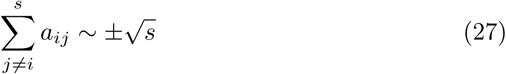

By assumption, this sum is an unbiased random walk of mean 0 and typical step size 1; hence, after *s* steps (with *s* large enough), no matter the distribution of *a_ij_*, the random walk will typically have gone a distance of order 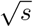 toward either positive or negative values. There are two important assumptions here: one of large numbers, which can only be checked numerically (in fact, we can often find good agreement even for *s* ~ 10), and one of randomness which is discussed more carefully in Sec. 5. If one accepts these assumptions for now, the final interaction strength perceived by any species falls roughly in the range

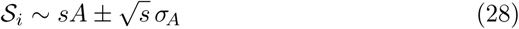

This is a crucial point: if we consider different systems with the same values of *A* and *σ_A_*, but different values of *S* or *c,* then we can see from (28) that, the larger and more connected the pool, the smaller the relative variation in 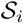 compared to its mean. In fact, for very large communities, there will be practically no difference in the interaction strength perceived by different species, making them *mean-field* (all species are effectively identical from the point of view of interactions) rather than disordered.

Instead, two pools of different size *S* have “equal” interaction strength and heterogeneity if their pools exhibit the same values of

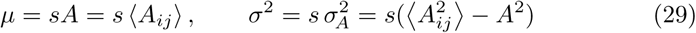

rather than the same values of *A* and *σ_A_*. To understand this, we can see that, no matter the pool size or connectivity, a species’ total interaction with others is given by

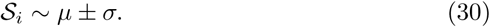

We can indeed show that *μ* and *σ* (in addition to a few other parameters) are what controls the properties of the assembled state – systems with different *S* but the same *μ* and *σ* will obey the same trends in terms of diversity, stability, abundance distribution, and so on. This rescaling means that systems with more connections between species (i.e. larger *S* or larger *c*) must have proportionally more varied interactions to retain the same *σ*, so that heterogeneity still matters and is not “averaged out”.

#### 4.1.3 Tradeoffs and global stability in large systems

We now further investigate this inverse perspective: instead of asking how *community parameters μ* and *σ* change with diversity *S* if we keep the same distribution of *individual parameters A_ij_*, we ask how these *A_ij_* should change with *S* to achieve a given *μ* and *σ.* This simple question actually contains an important insight on the properties of large assembled communities. It is easy to see that if one wants to fix connectivity *c* as well as *μ* and *σ*, but increase *S*, then necessarily interactions should vary like

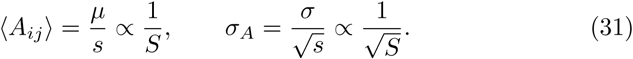

For large enough systems *S* ≫ 1, if we maintain the same values of *μ* and *σ*, we thus expect *σ_A_* ≫ 〈*A_ij_*〉. Having a standard deviation much larger than the mean is not achievable with “reasonable” distributions^7^ unless we allow the interactions to take any sign (the mean can thus go to zero while the variance remains large).

Hence, due to simple mathematical constraints, large systems must have at least one of these four properties: large *μ*, small *σ*, low connectivity *c*, or mixed interactions allowing for all signs. For instance, a large competitive community where all species can potentially interact (*c* = 1) must have either large *μ*, meaning very strong competition and low biomass, or small *σ*, meaning that heterogeneities in species interactions cease to matter and the system is effectively mean-field. This “biomass–heterogeneity” tradeoff in large competitive communities, which we emphasize comes from simple algebra independent of any model, is indeed very apparent in our analysis of resource consumer models in [4].

This tradeoff has important consequences: as shown in recent work [40], small *σ* is a condition under which a complex ecosystem might be reduced to a single dimension, leading to predictions of “universal” behavior [9]. Furthermore, small *σ* and small *ζ* taken together constitute the basic hypothesis of neutral theory – that we can take all species as identical in large communities, despite their very real heterogeneities. Hence, the fact that simple algebraic constraints could limit *σ* in large competitive communities provides some formal grounding for the intuitions behind “universal” and neutral dynamics.

### 4.2 Couplings rather than interactions

#### 4.2.1 Allowing heterogeneous self-interactions

At multiple points in the discussion above, we have suggested that interactions are not, by themselves, the quantities that control the equilibria: instead, that role is taken by *couplings,* i.e. the ratios of interactions to self-interaction, which represents how much the influence of another species could cause one to deviate from its independent logistic trajectory.

In Sec. 2, we presented results assuming that all self-interactions were equal, *D_i_* = 1. But in fact, we can easily see that we can transfer our conclusions to the Lotka-Volterra equilibrium equation (3) *divided by D_i_*^8^:

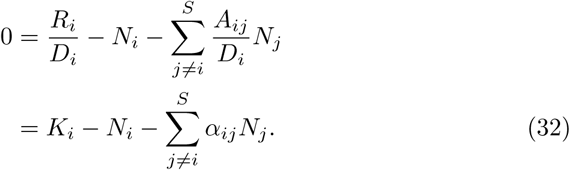

Here, we recall *K_i_* = *R_i_*/*D_i_* the carrying capacity of species *i,* and *α_ij_* = *A_ij_ /D_i_* the coupling of species *i* to species *j.* Thus, all the results we presented in Sec. 2 still hold if, instead of using the variance of growth rates, we use that of carrying capacities, and instead of using the statistics of *A_ij_*, we use those of *α_ij_*. In other words, we formally redefine the parameters of (4) as follows:

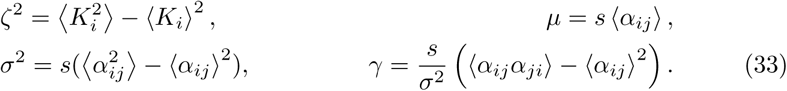

We can relabel the parameters and retain the same mathematical structure, and their interpretation does not change too dramatically – except in the important case of *γ*, as we soon discuss. Previously in this section, we used *A* = 〈*A_ij_*〉 and *σ_A_* extensively to discuss simple and general properties of the dynamics – the distribution of the interactions *A_ij_* is otherwise entirely unspecified, and could take arbitrary forms (Gaussian, exponential, uniform, gamma, multimodal …) except for the pathological. We saw in examples that almost all other properties of this distribution *P*(*A_ij_*) were irrelevant to assembly for large enough communities, one of the very counter-intuitive and attractive traits of disordered systems. In fact, we see now that, if self-interactions differ, we should use the *α_ij_* instead; but thankfully, results remain insensitive to the details of *P*(*α_ij_*). The subtlety is found in the fact that *σ* now includes not only the heterogeneity of interactions, but also that of self-interaction – in other words, species with identical interactions but very different growth rates and carrying capacities will have large *σ*, with all the consequences shown in Sec. 2 in terms of coexistence and multistability.

This of course requires that *D_i_* ≠ 0 – and in fact that *A < D_i_*, as the case of very strong interactions *A > D_i_* tends to lead to very unstable dynamics that we wish to avoid entirely in this approach^9^, as discussed in 1.2.1.

#### 4.2.2 Reinterpreting *γ*

However, there is one crucial change in interpretation. The parameter *γ* ∊ [−1, 1] represents the average degree of reciprocity between the dynamical coupling of species *i* to *j*, and that of *j* to *i*. The extreme case *γ* = 1 indicates fully symmetrical couplings *α_ij_* = *α_ji_*, while *γ* = −1 means anticorrelated couplings. Unfortunately, while this parameter is the one that truly encapsulates how asymmetry between interaction partners affects the assembly process, its interpretation is very subtle and counter-intuitive.

Let us first discuss *γ* in the previous case where all self-interactions are equal, *D_i_* = 1, as even that simpler situation has some difficulties. If *A* = 0, then *γ <* 0 suggests trophic interactions, since *A_ij_* and *A_ji_* will typically be of opposite signs. However, if A ≫ *σ_A_*, then even with *γ* = −1, *A_ij_* ~ *A* + *σ_A_* and *A_ji_* ~ *A* − *σ_A_* will have the same sign. They will simply be maximally anticorrelated: for every pair of species, one will affect the other much more than it is affected by it. This can for instance give rise to “rock-paper-scissors” intransitive competition.

If we now return to varied self-interactions *D_i_* ≠ 1 so that couplings are different from bare interactions, even that refined notion of reciprocity becomes more complex. Consider for example the classic resource competition models [21, 10, 39]. By construction, interactions come from two species harvesting the same resource and are therefore perfectly symmetrical, *A_ij_* = *A_ji_*. However, couplings are *not* – in fact, depending on parameters, the model can give any *γ* between −1 and 1, and most plausible parameter values correspond to *γ <* 0. In any pair of competitors, each reduces the other’s growth equally, yet this will on average have a much larger impact on the dynamical trajectory of one species than the other.

Hence, the proper way to understand *γ* is in relative terms: no matter whether the interaction is positive or negative or even symmetrical, if *γ* = −1, one species will respond much more to the interaction (in the sense that it will be more affected compared to the trajectory it would have if alone), while for *γ* = 1 they will respond equally. Given that assembly is a process of selecting species that are most compatible among a larger pool, it can be understood that what really matters is not the net outcome of each interaction, but how differently it affects the partners’ dynamics. But despite its predictive power, any intuitive interpretation of this parameter can only be made in the context of a more concrete model.

### 4.3 Computing the assembled state

At the start of Sec. 4, we showed that the equilibrium of the Lotka-Volterra equations (3), which we recall is given by

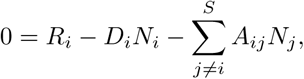

can be rewritten in all generality as

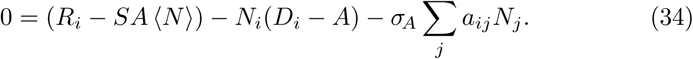

given the mean *A* = 〈*A_ij_*〉 and the variance of interactions 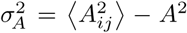, with the rescaled interactions *a_ij_* = (*A_ij_* − *A*)/*σ_A_*.

As we argued in Sec. 4.2.1, the quantities that truly control the dynamics are not “bare” interactions *A_ij_*, but couplings

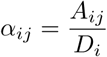

which represent how much species *j* can cause species *i* to deviate from its baseline (logistic) trajectory controlled only by self-interaction *D_i_*. We now rewrite

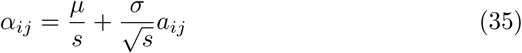

with

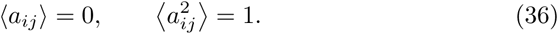

Then, we have

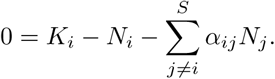

and instead of (34) we get

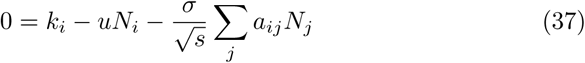

where we have defined the effective self-interaction

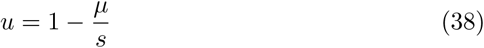

and the effective carrying capacities

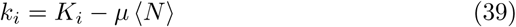

Following previous work [6], we use the cavity method from physics [29] to find the properties of the equilibria of equation (37). Technical details are given in Sec. 5. To understand the results and their limitations, it is enough to grasp the following simplified discussion, introducing some important concepts.

Let us take an assembled state created from a pool with *S* − 1 species. When the final species *i* invades, if *S* is large, the properties of the assembled state should not change significantly^10^. Hence, we imagine that we can move from *S* − 1 to *S* species holding everything constant, except for the fact that the other species have their final abundance *N_j_* shifted a small amount, in the direction suggested by their couplings with *i*,

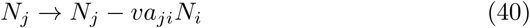

with *v* a positive “response coefficient” that we can compute. Inserting this shifted abundance in (37), a feedback is created on species *N_i_*. This is where the parameter *γ* enters the picture: if *γ >* 0 the feedback is positive (e.g. a species reduces its competitors’ abundance, which in turn allows it to grow more), while if *γ <* 0 the feedback is negative (e.g. a predator reduces its preys’ abundance, which stifles its growth). This explains how negative *γ* can play a stabilizing role in the dynamics, as we have seen above starting in Sec. 3.3.

The finite trick is to see that this equation expresses *N_i_* as a random variable obeying:

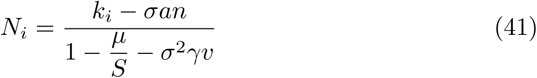

where *a* is a Gaussian random variable of mean 0, derived from the random walk (27) on the *a_ij_*’s, and *n* is a random variable drawn from the distribution *P*(*N*) of surviving species. This simplification can be made technically precise, and relies on a crucial property of disordered systems: there is no correlation between the *a_ij_*, which represent direct couplings, and the *N_j_ before* the shift (40), which are shaped by all the other indirect couplings in the system. These two could not be taken as independent parameters if each species depended strongly on a few others, but the disordered nature of interactions means that species abundances are shaped mostly by collective dynamics involving the whole community. A failure of this approach to correctly predict the behavior of some model could signal that local ordered dynamics prevail over collective disordered dynamics.

The second central assumption that is made here is the existence of a single global attractor, meaning that no matter which initial condition is taken, the same assembled state is reached provided all species are given a chance to invade when they can. This premise is hidden in the fact that we assume that equilibrium species abundances do not shift drastically with the addition of a new species. As we have mentioned in Sec. 3.3, we can compute a quantity that becomes infinite precisely at parameter values for which there ceases to be a single attractor, thus signalling the appearance of potential tipping points. Methods in physics have been derived for the express purpose of investigating such multistable regimes, and quantifying the number and stability of attractors. In the case of Lotka-Volterra dynamics, however, this task is ongoing.

The assembled state must be feasible and uninvadable, and therefore, studying it reveals what controls species coexistence. But it also contains information on the ecosystem’s sensitivity to external change: the quantities that arise naturally when computing the assembled state are long-term responses to a press perturbation, i.e. how the whole community’s equilibrium shifts in response to some species being pushed toward an abnormal abundance. This can come from durable environmental changes that affect the fitness of some species, but also from the introduction of a new invader into the system.

## 5 Detailed calculations

Readers who are only interested in the general spirit of the method can stop here, as we turn to the practicalities of using it and the dynamical intuitions it brings. In this section, we follow in closer detail the derivation of the equilibrium solution for a large Lotka-Volterra system first presented in [7]. For the mathematically minded reader, let us note that related calculations, using more technical methods from disordered systems, had previously been achieved for replicator equations (which are closely related to Lotka-Volterra) and mostly discussed within physics [32, 41]. Earlier results relied on the “replica method”, limited to symmetrical interactions (hinging on the fact that there then exists a Lyapunov function, as noted long ago by MacArthur for resource competition [21]). Later results employed a “dynamical generating functional method” which allows to find not only equilibria but a single equation for the effective dynamics of the entire system [32], although that equation is partially implicit and can only be solved by ansatz.

Bunin [7] proposed the first derivation for the equilibria of Lotka-Volterra systems, using a technically simple implementation of the “cavity method” [27]. This calculation is comparatively straightforward while retaining many important intuitions of disordered systems, and its application to ecological models is immediate, which makes it especially valuable for introductory purposes.

### 5.1 The cavity trick

The assembly process shapes the community by changing the abundances of species until an equilibrium is reached. From the imposed properties of the pool, assembly uses the abundances as its tool to select and reweigh species features, crafting a state that is feasible, stable and uninvadable by any other species in the pool. Hence, all the properties of the solution can be derived from the distribution of abundances and their correlations with species features (such as carrying capacities and interactions).

The logic of the cavity method is remarkably simple: 1) we compute what happens after a single invasion in an already large community, and 2) we then require that the equilibrium features before and after the invasion be unchanged. The second part means that we are computing the fixed point of the invasion process, i.e. an equilibrium which cannot be invaded nor destabilized, rather than a single-step invasion as is often done in population genetics (testing whether a given mutant species can invade).

As shown in Fig. 6, the calculation of the invasion is made simpler by assuming that, even though the invader itself retains its nonlinear dynamics, the effect of the invader on each already present species is small enough to be approximated by a *linear* change of their abundance. This assumption may seem intuitive if the final abundance of each species is the result of many interactions, so adding one more should not cause a dramatic change. A more formal justification is given in Appendix A.

**Figure 6:**
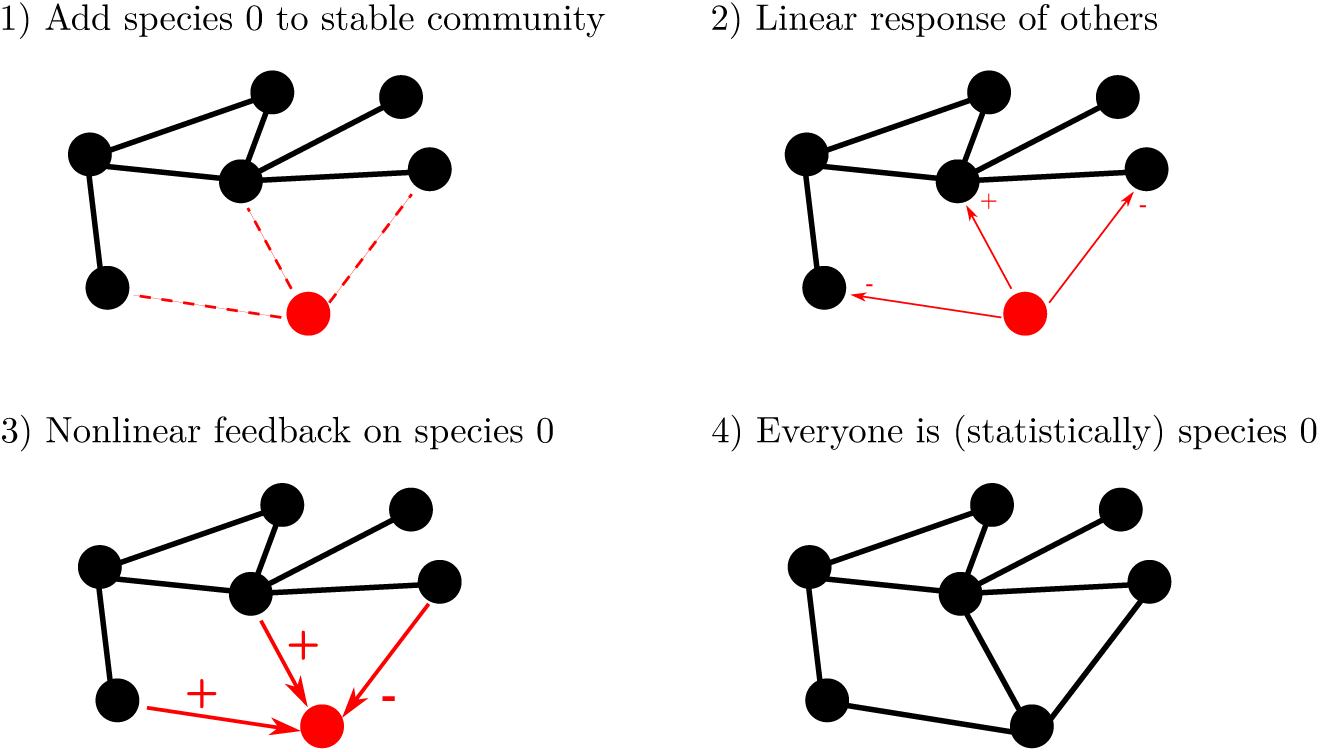
Cartoon of the cavity method.

Let us introduce species 0 into a community of *S* − 1 species. Its equilibrium abundance *N*_0_ changes other *N_i_* a little, as they now obey:

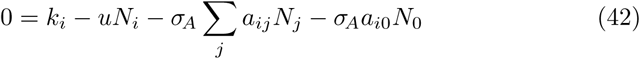

(where we recall that 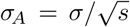). Formally, this looks like changing their carrying capacities *k_i_* a little

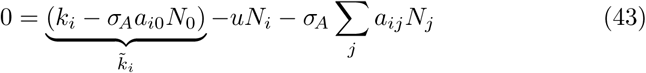

where 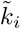 is a slightly displaced carrying capacity. We now assume that *N_i_* responds linearly to this small perturbation: using a Taylor expansion around the previous equilibrium abundance 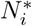, calculated without species 0, we can write

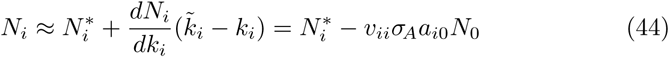

where we define the response coefficient,

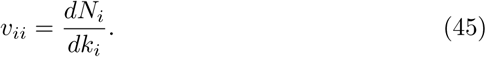

This seems unhelpful at first, since *v_ii_* is just another unknown in the problem, but we will show that creating this new set of unknowns actually helps us further down. To interpret *v*, let us remark that, if species were independent, then

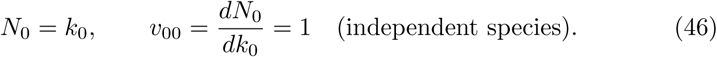

Hence, *v* ≠ 1 indicates that species abundances are not defined solely by their intrinsic fitness, but by their effective fitness including interactions – it is possible to change their carrying capacity without affecting their abundance proportionally.

In fact, all other species in the community perceive the effect of adding species 0, so a better approximation of the full linear response would include “cross-terms” i.e.

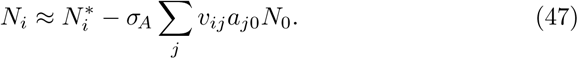

with

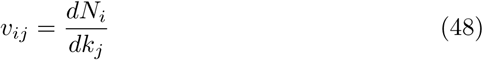

the long-term response of species *i* to a change in the carrying capacity of species *j*, including all the indirect effects from other species. In fact we show later that these indirect effects tend to cancel out and disappear, but let us include them for now, for the sake of completeness.

Now, this shift in others’ abundance feeds back into the equation for *N*_0_, which tells us how it depends on 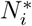 the abundances before its invasion

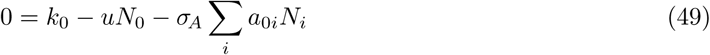

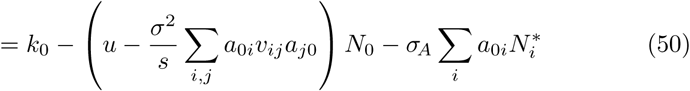

The issue is that it contains a double sum over all the unknown *v_ij_*, representing the overall feedback of species 0 on itself through the rest of the community. A crucial simplification is that, due to the disorder of interactions *a_ij_*, these many unknowns collapse into a single one that can easily be determined later: this large sum tends to an average value, as we explain very roughly here and further detail in Appendix A. Since 〈*a*_0*i*_〉 = (*a*_*j*0_) = 0, all terms that have *i* ≠ *j* are fluctuations around 0, while each term 〈*a*_*i*0_*a*_0*i*_〉 = *γ* has a significant contribution to the sum. Hence, as we noted before, the main contribution comes from the reponse of *N_i_* to a change of *k_i_*, rather than from cross-terms:

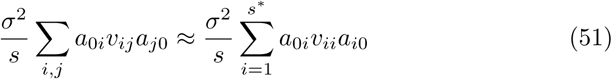

where *s\** = *sϕ* is the number of species that interact with species 0 and are still alive in the assembled state (we recall that *ϕ* is the overall fraction of surviving species). This sum is an average over all the interaction partners of species 0, and it is in fact “self-averaging”. This is physics jargon to say that, given a large number of species to interact with, it tends to become a single well-defined number for all species in the community, with fluctuations that become small as *s* becomes large,

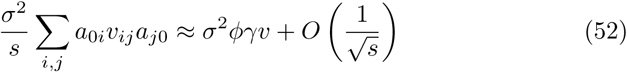

where

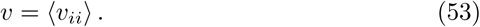

Finally, we obtain the effective equation for *N*_0_ which will be solved below.

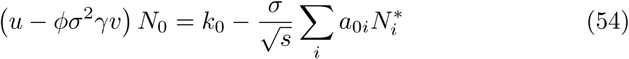

### 5.2 Universality of cavity results

Isolating *N*_0_ on the left-hand side,

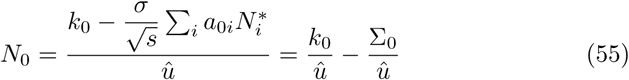

where

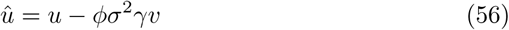

and we have defined

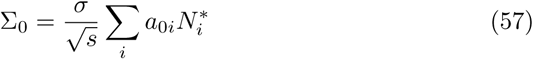

We can see in *N*_0_ two main contributions (stretched by a factor 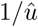 that is common to all species in the community): the effective carrying capacity *k*_0_, and a fitness shift Σ_0_ arising only from to the *heterogeneity* of interspecies interactions. The former is model-dependent: we have no reason to constrain the distribution of carrying capacities *K*_0_ which is a property of the species pool (we recall that *k*_0_ = *K*_0_ − *μ* 〈*N*〉). But the latter becomes universal in the limit of many interactions. This relies on the sum 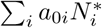 being effectively a random walk.

Here, we are going to assume that interactions between species are not correlated *in the pool* (we discuss a weaker assumption that is still tractable in Sec. 7.4). This lack of correlation has two consequences. First, note that *a*_0*i*_. is uncorrelated with 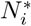: the abundance of species *i before species* 0 *was introduced* depends on *a_ji_* for *j* ≠ 0, but bears no relationship to *a*_0*i*_. More importantly, *a*_0*i*_ and *a*_0*j*_ are uncorrelated, which means that all terms in the sum above are uncorrelated variables, and if they have finite variance, then the sum must tend to a Gaussian law by the Central Limit Theorem. This Gaussian law has mean 0 and variance *sϕ*〈*N* ^2^〉^*^, where the mean is taken here over *P*(*N\**) the equilibrium distribution of values for 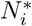.

Hence, for any model where interactions have finite variance, the final abundance distribution will be composed from the model-specific distribution of carrying capacities, and from a generic Gaussian term emerging from interactions^11^. And even if *P*(*N\**) had a model-specific shape due to *P*(*K*), the interaction term, being Gaussian, is always entirely parametrized by the first two moments of *P*(*N\**), meaning that one never needs a lot of information on *P*(*K*) to correctly evaluate the effect of interactions.

This entire reasoning however weakens if *P*(*K*) – and therefore *P*(*N\**) – have fat tails; any other structure (e.g. multimodal *P*(*K*)) will somewhat increase the necessary number of species and interactions to converge toward the Gaussian distribution of *σ*_0_, but fat tails may prevent this convergence altogether (if variance is formally infinite) or make it dramatically slow. This possibility would call for an extreme value calculation instead: what would matter in Σ_0_ would not be its variance anymore, but its extreme values, which obey a different but still universal distribution.

We will now assume that *P*(*K*) is “well-behaved” (e.g. Gaussian or uniform) so that it is enough to know its mean and variance to derive *P*(*N\**). We discuss in Sec. 6 the sensitivty of results to choices of *P*(*K*).

### 5.3 Abundance distribution

Let us recall equation (55) which defines the equilibrium abundance *N*_0_ of a representative species in the community as a sum of two random contributions:

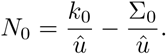

This expression allows for *N*_0_ *<* 0, hence, *N*_0_ is a “virtual abundance” that signals that some species could invade and attain abundance *N*_0_ if *N*_0_ *>* 0, or that the species could not invade durably if *N*_0_ ≤ 0. (Note that the more traditional approach to assembly is to check whether a species can grow immediately at the start of its invasion, without predicting the rest of its trajectory; here on the contrary we check whether it could remain at infinitely long times, after all interactions with the rest of the community have played out).

This virtual abundance has been computed from the carrying capacities of invaders *P*(*K*_0_) and the abundances of surviving species *P*(*N^*^*). This means that, if species 0 itself survives, it becomes representative of survivors. If we can compute the probability distribution *P*_0_(*N*_0_), then *P*(*N*\*), must be the same distribution restricted to positive values.

An important conceptual trick in this approach is that, instead of computing species abundances for a given community, i.e. a fixed choice of interactions and carrying capacities, we suppose here that all abundances are fixed (even if we do not know their values yet) *except* for that of species 0, and try to compute the typical distribution of *N*_0_ over the whole range of possibilities for *a*_0*i*_*, a_i_*_0_ and *K*_0_. From it we will compute the “typical” abundance distribution *P*(*N^*^*), i.e. the one that we will see on average over all possible communities whose parameters are drawn from these probability distributions.

Since we know *N*_0_ as a function of 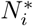 in a given system, and *P*(*N^*^*) as a function of the distribution *P*_0_(*N*_0_) over all possible choices for interactions and carrying capacities, we have a closed system that can be solved through the method of moments. First, we compute the first two moments of *P*_0_(*N*_0_) as a function of those of *P*(*N^*^*). We can take the mean over *P*(*a*_0*i*_) and *P*(*K*_0_) of both terms in (55), we find

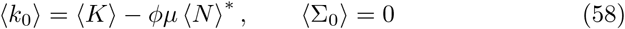

where we used (*N*) = *ϕ*〈*N*〉^*^ (since when we introduced 〈*N*〉 earlier, we took the mean over all *S* species, including those that are dead), and the facts that 〈*a*_0*j*_*N_j_*〉 = 〈*a*_0*j*_〉 〈*N*〉 and 〈*a*_0*j*_〉 = 0. Hence

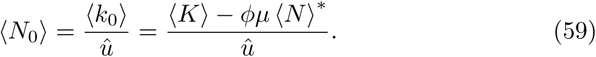

For variances, we can similarly find

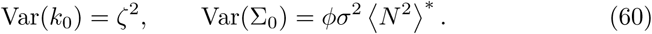

Hence, if *k*_0_ and Σ_0_ are independent variables (see below otherwise) the variance of the sum is the sum of variances and

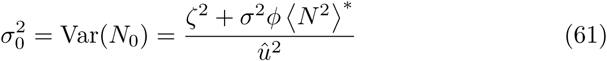

Following our assumption that *P*(*K*) is well-behaved, these two moments are enough and we approximate *P*_0_(*N*_0_) by a Gaussian law with the mean and variance just given,

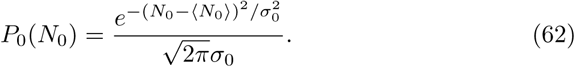

From there, we can give a definition to 〈*N*〉*\**, 〈*N*^2^〉^*^ and *ϕ* by restricting *P*_0_ (*N*_0_) to its positive values

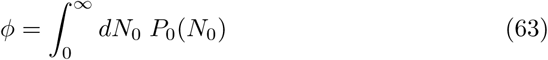

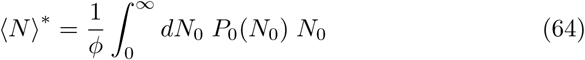

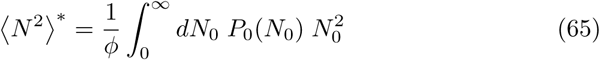

where the 1/*ϕ* factor ensures normalization. On the other hand, *v* is given by taking the derivative (48) in (55)

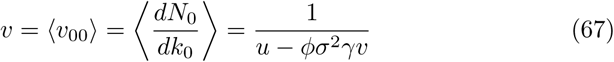

which is thus coupled to the rest through *ϕ* (and by appearing in *P*_0_). We now have a closed set of four equations for four unknowns: *v*, *ϕ*, 〈*N*〉*\**, 〈*N*^2^〉* which appear on the left-hand side of each equation and in *P*_0_ (*N*_0_). These equations can be solved together numerically. It is then immediate to compute quantities such as total biomass

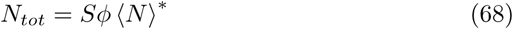

and Simpson index

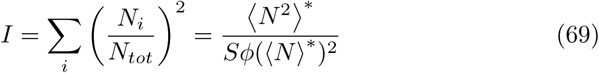

#### 5.3.1 Correlations between *K_i_* and *A_ij_*

Equation (61) assumed *k*_0_ and Σ_0_, and hence *K_i_* and *A_ij_*, to be independent variables. If there is a correlation between them, then

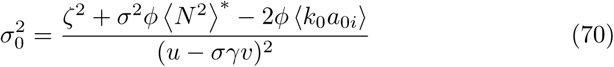

where

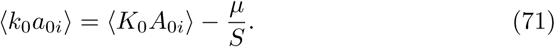

Hence, such correlations can be accounted for, at the cost of adding a new parameter. Other simple extensions are discussed in Sec. 7.

#### 5.3.2 Relationship to empirics

The fact that the theoretical abundance distribution is (approximately or exactly) a truncated Gaussian contradicts much evidence on empirical abundance distributions being fat-tailed – possibly log-normal or even power-law. No fault lies with the mathematical analysis, as it is in perfect quantitative agreement with many simulation models, see [4]. Hence the issue will occur for any system with Lotka-Volterra dynamics (1) with random interactions and carrying capacities drawn from many usual distributions, and without noise.

There are at least three solutions to this problem. The first is a wide distribution of carrying capacities: let us recall that in Sec. 2.2.3, we noted that many competitive models [17, 26] assume^12^ that interactions scale like

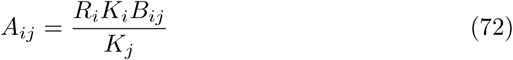

with some narrowly distributed *B_ij_* (e.g. uniform or Gaussian). In this case, we noted that we could write

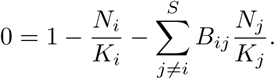

Hence, by having *μ, σ* and *γ* be the properties of the *B_ij_*, the results above predict that the distribution *P*(*N_i_*/*K_i_*) is a truncated Gaussian. If *K_i_* follows a lognormal distribution for instance, as seen in plant monocultures [34], then *N_i_* at equilibrium will have an even wider distribution.

This explanation however assumes that the wide spread of species abundances is due to some intrinsic species properties, which define *K_i_* in a given environment. Under this explanatory scenario, our analysis is compatible with fat-tailed abundance distributions, but does not explain their emergence. However, there are at least two ways in which the dynamics themselves could produce fatter tails: first, the addition of noise, which is a central ingredient in neutral models that successfully predict abundance distributions. Second, the role of spatial aggregation in the data, as it is well known that aggregating over Gaussian distributions with different variances can create the appearance of more complex distributions and even fat tails - indeed, this is seen in an extension of this work to metacommunities (to be published).

### 5.4 Selection by assembly: the case of carrying capacities

Starting from this section, the core of the method is established and we simply show how various quantities of interest can be calculated using the results above.

We have seen above that equilibrium abundances are made up of two independent contributions: one coming from the effective carrying capacity, the other a shift in abundance due to the heterogeneity of interactions

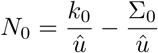

We also recall that

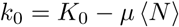

If we want to know the equilibrium distribution of carrying capacities 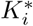, we start from the joint distribution of abundance *N*_0_ and effective carrying capacity *k*_0_, which we will then restrict only to surviving species *N*_0_ *>* 0.

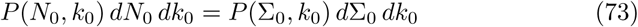

hence

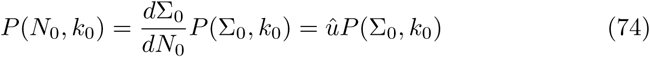

Now, since Σ_0_ and *k*_0_ are independent, *P*(Σ_0_, *k*_0_) = *P*(Σ_0_)*P*(*k*_0_), so

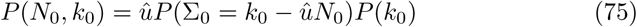

We argued above that *P*(Σ_0_) is Gaussian with mean 0 and variance 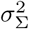

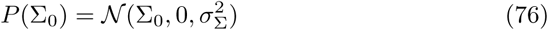

with

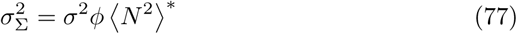

Now, we want to isolate *P*_0_ (*N*_0_) within *P*(*N*_0_, *k*_0_), so that the integral over *N*_0_ *>* 0 causes the moments we have already computed, *ϕ*, 〈*N*〉 and 〈*N* ^2^〉, to appear. We recall

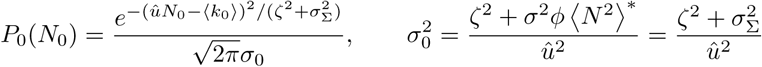

With some manipulations (assuming a Gaussian distribution for *K_i_*), we can see that

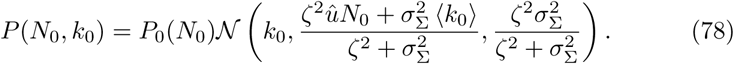

In particular, we find the averages over surviving species

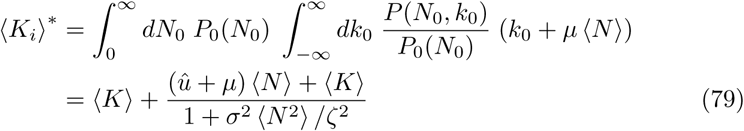

and

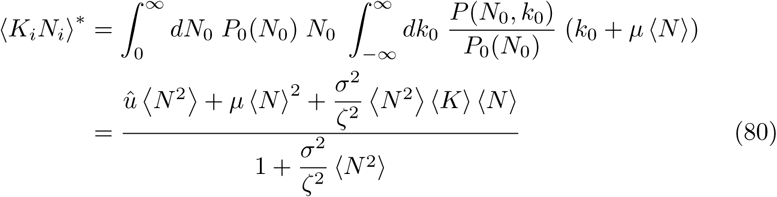

where we recall 〈*N*〉 = *ϕ* 〈*N*〉*^*^*, 〈*N*^2^〉 = *ϕ* 〈*N*^2^〉*^*^*.

### 5.5 Productivity

Given the abundance distribution *P*(*N_i_*), it is possible to compute the productivity distribution – or at least the distribution of growth on external resources,

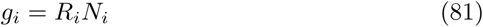

which acts as a proxy for productivity. If the quantities that are drawn independently are *K_i_*, *α_ij_* and *D_i_*, then straightforwardly

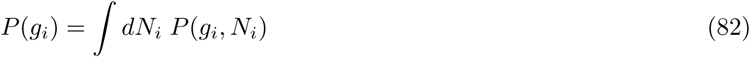

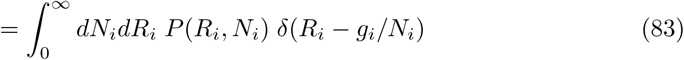

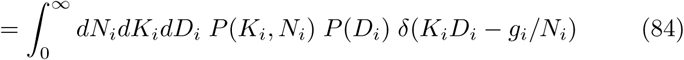

and we get *P*(*K_i_*, *N_i_*) from the previous section. In particular, taking the average over surviving species, we find (if all *K_i_ >* 0)

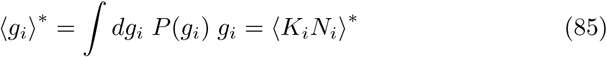

where we used 〈*D_i_*〉 = 1. If however the independent quantities are *R_i_*, *A_ij_* and *D_i_*, the calculation is less straightforward: we must now start from

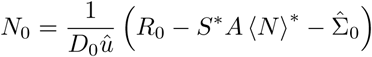

where

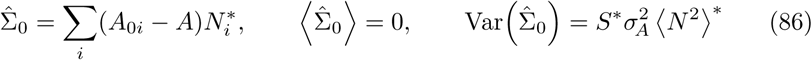

and we now have 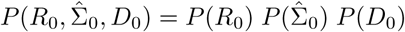, hence

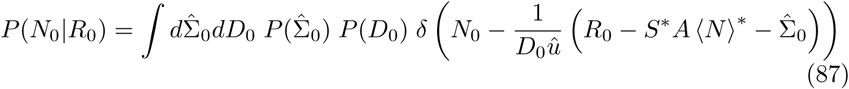

and we get the total productivity

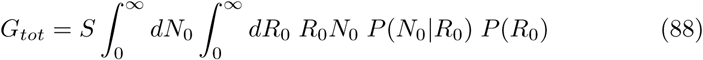

### 5.6 Response to perturbations

Let us now assume that some small permanent (press) perturbation *ξ_i_* is added to the system at equilibrium (acting only on the surviving species)

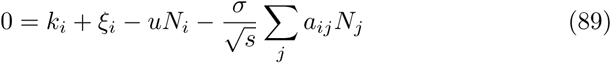

We can write 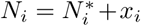, with *x_i_* the displacement around equilibrium, which obeys

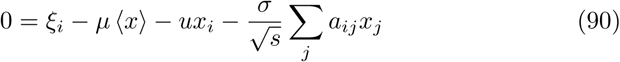

a very similar equation to that obeyed by equilibrium abundances, except for the fact that the carrying capacity *K_i_* has been replaced by the perturbation *ξ_i_*, and of course that *x_i_* is not constrained to be positive. Following the same logic as before, we isolate the response of species 0

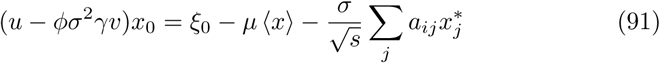

If there are no constraints on *x*_0_ (see Sec. 5.6.2 below), we can simply assume

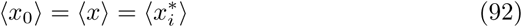

and thus get the average displacement:

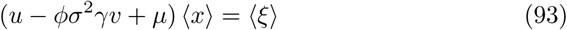

#### 5.6.1 Multistability order parameter

To test whether the equilibrium we are obtaining as the fixed point of the invasion process is globally stable, or might be exited through some small perturbation in the appropriate direction, we must consider the response to a random perturbation with zero mean,

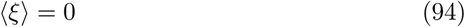

representing random shifts in relative fitness that advantage some species over others, without changing the mean species fitness. Since the mean is zero, the response to the perturbation is characterized by the variance,

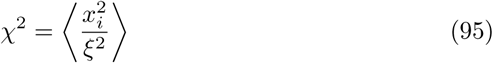

Now the equation above, using 〈*x*〉 = 0, can be taken to the square to get:

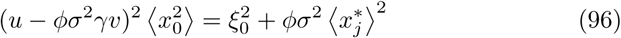

hence

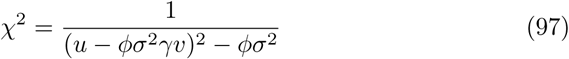

which is seen to diverge when *σ* = *σ_c_* such that

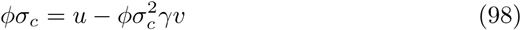

and by using (*u* − *φσ* ^2^*γv*)*v* = 1 we find as above in (15)

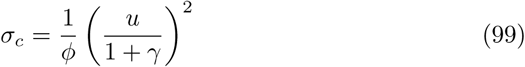

#### 5.6.2 Secondary extinctions

In the case of secondary extinctions following the extinction of species 0, the perturbation is

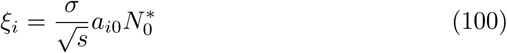

Now however, the perturbation is potentially large enough that we may need to impose 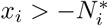, meaning that the response cannot cause a species’ abundance to become negative. Hence, in a way similar to our previous calculation on abundances, the average change in abundance due to the extinction is

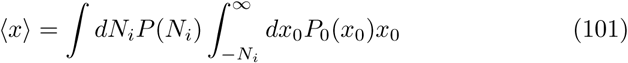

while the fraction of secondary extinctions is given by

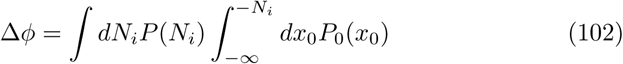

#### 5.6.3 Variability

Calculations for the variability are more technical and left in Appendix.

## 6 Sensitivity testing

### 6.1 Distributions

In this section, we test how sensitive the results above are to the choice of distributions for growth rates and interactions. Throughout, we have assumed that these distributions had finite mean and variance, which is not the case for, e.g., a power-law tailed distribution P(x) *~ x*^−*a*^ with exponent *a ≤* 3.

For reasons explained in Sec. 5.2 above, we expect results to be more sensitive to the distribution of growth rates than to that of interactions. Indeed, Fig. 7 shows that, while Gaussian or uniform distributions lead to very good agreement with predictions, distributions with fatter tails (e.g. exponential tails) – and a fortiori multimodal distributions – lead to significant deviations from our baseline predictions. It is however not difficult to adapt the theory to fit any distribution *P*(*R_i_*) as long as it is known.

**Figure 7:**
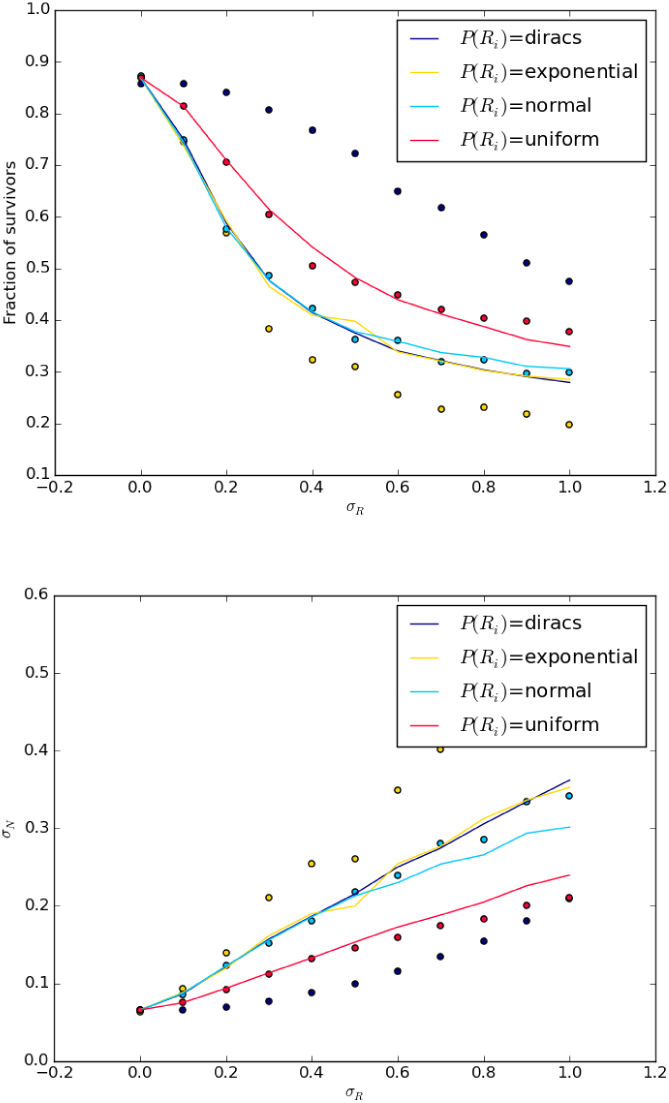
Sensitivity of results to distribution of growth rates *P*(*R_i_*).

### 6.2 Interaction strength

We now test how small the interaction strength must be compared to self-interaction for cavity predictions to work. Fig. 8 shows that when average coupling strength is significantly larger than 0.1, the method stops being as accurate, in particular with fewer species surviving than expected.

**Figure 8:**
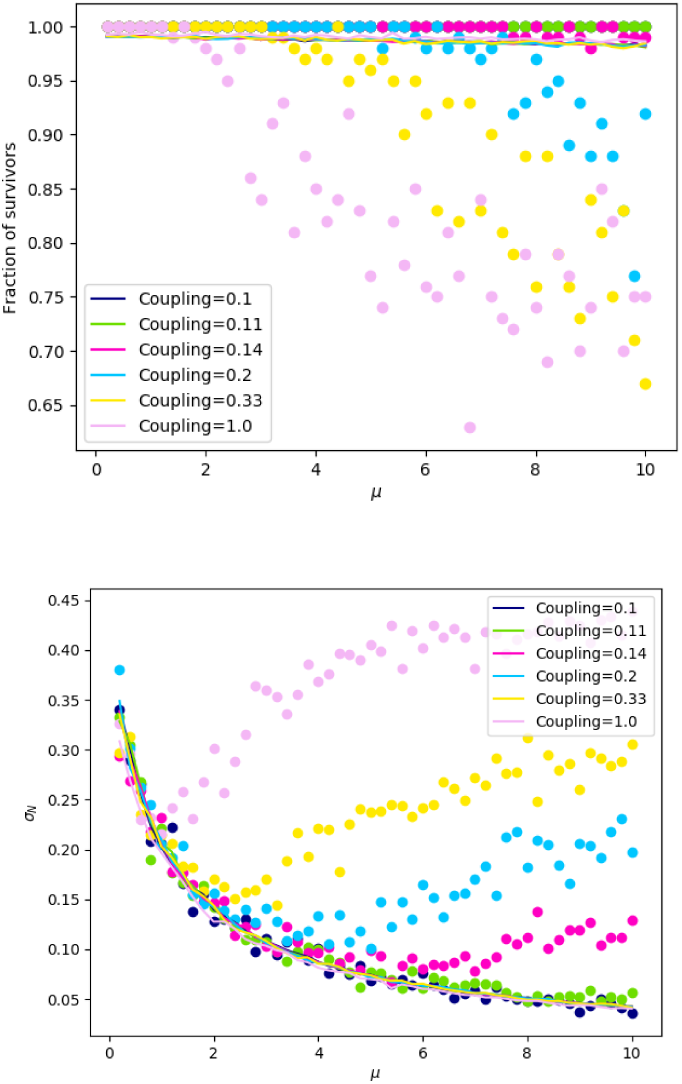
Sensitivity of results to interaction strength. By varying the connectivity from 0.1 to 1, we change the average interaction strength needed to achieve the same value of *μ.* Each curve is labelled here by the average coupling strength needed to obtain *μ* = 10.

## 7 Extending the model

There are two main types of limitations to the model so far. The first is dynamical: we started from Lotka-Volterra dynamics, which might work as an approximation in many limits, but do not always enable us to find the equilibria reached by more complex dynamics. Since the most usual variant to Lotka-Volterra dynamics is having a nonlinear functional response, especially a saturating one, we treat that case below to show that it is still largely amenable (at least approximately or qualitatively) to our approach.

The second limitation is structural. Even though species are heterogeneous, we have treated them as if they all belong to the same functional group: their traits are all drawn from the same distributions. These distributions may be multimodal, but that does not convey the fact that different positions in the community may be truly different in many respects at once. There are two simple ways to address this concern, one discrete and one continuous.

The continuous approximation, which we call “ordered species”, supposes that we can characterize the network by a simple axis of possible niches or roles, knowing only how various traits (e.g. carrying capacity, interaction mean and variance) correlate to linear order with position on that axis.

The discrete approximation, which we call “species groups”, instead divides the network into distinct groups, each disordered inside, but with deterministic patterns within and between groups. In essence, this is extending simple reductionist descriptions (e.g. one predator and one prey) by replacing each species by a group of heterogeneous but functionally similar species, treated like a disordered community.

### 7.1 Functional response

The coefficients *A_ij_* come from two distinct factors: first, how much an *individual* of species *i* can affect an individual of species *j*, which may be a property of their behavior and biology, and second, how often these individuals may meet and interact. For simplicity, let us say that the individual-level effects are fixed. Then, the frequency of pair encounters is what controls interaction strength. The interaction between species *i* and *j* is independent from any other interaction in relatively “sparse” environments where any pair can meet, and adding new individuals or new species always increases the number of such meetings. On the other hand, interactions may saturate if there is a limiting factor such as plants competing only with their neighbors, or a predator taking time to handle prey, preventing it from consuming more than a given number of prey among those available in a given timespan.

The same reasoning led Holling [14] to postulate a saturating functional response in the interaction between two species: instead of being a number, *A_ij_* becomes a function of abundances such as

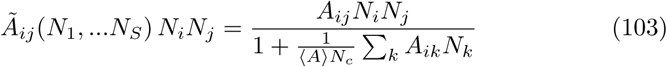

with constants *A_ij_*, their average 〈*A*〉 and the population threshold *N_c_*. This means that the interaction between species *i* and *j* cannot keep growing indefinitely with *N_i_N_j_* the product of their abundances, as in Lotka-Volterra equations. Hence, the sum is bounded

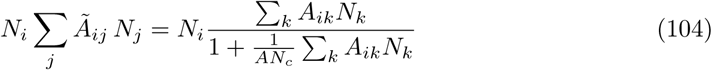

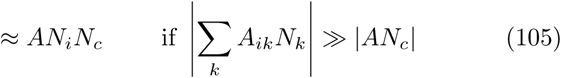

Analytically, it is possible to deal with the saturating functional response by a simple approximation. For each species, we replace it with a piecewise linear function:

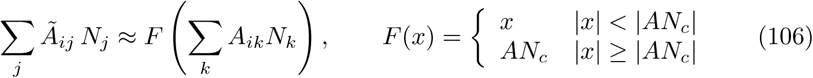

then at the community level, we use the real saturating function to evaluate the expected fraction *f* of species whose interactions are saturated.

For simplicity, we will “cheat” and instead apply the piecewise functional response to the *couplings* (which differ from interactions if species have heterogeneous self-interaction *D_i_*), i.e.

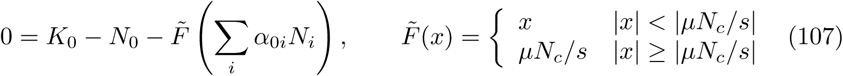

Avoiding this simplification is possible but requires keeping track of more parameters, and does not make much of a qualitative change.

Going through the same steps as before, then for each species

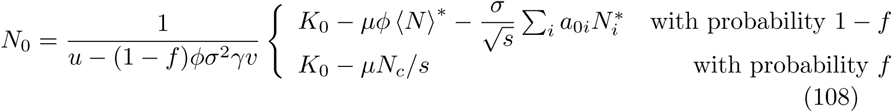

Notice the 1 ***–*** *f* factor in front of the feedback term in the denominator: only species whose interactions have not saturated will contribute to the feedback of the invader on itself through its interaction partners. The mean is now

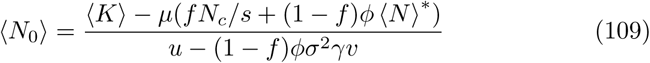

while taking the variance, we find

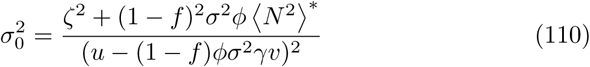

The rest of the calculations follow as in Sec. 5.3, except we additionally need to know the fraction *f* of species with saturated interactions. This is where the real saturation function reappears (slightly rearranged):

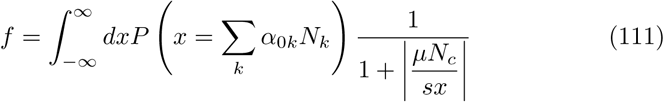

where, as usual, we can approximate *P*(*x*) by a Gaussian distribution with

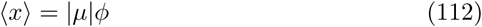

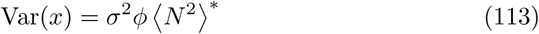

Note that if we want to make an equivalent model without saturation, we can apply the equivalence

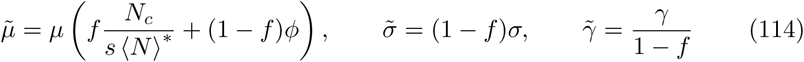

which is possible only if 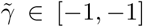. Then, a Lotka-Volterra system with parameters 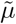, 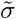 and 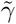 will have identical equilibrium properties to the system with a saturating functional response studied here. Of course, this equivalence can only be computed once we know *f* and 〈*N* 〉*, i.e. after having solved the calculation above for this system.

In Fig. 9, we show that predictions from our calculations match simulation results quite well, despite many simplifications which tend to create a more abrupt transition between the saturated and unsaturated regimes. By the same reasoning, we can easily extend the approach to any functional response, including an approximation of Type-III sigmoid response, with two thresholds instead of a single one.

**Figure 9:**
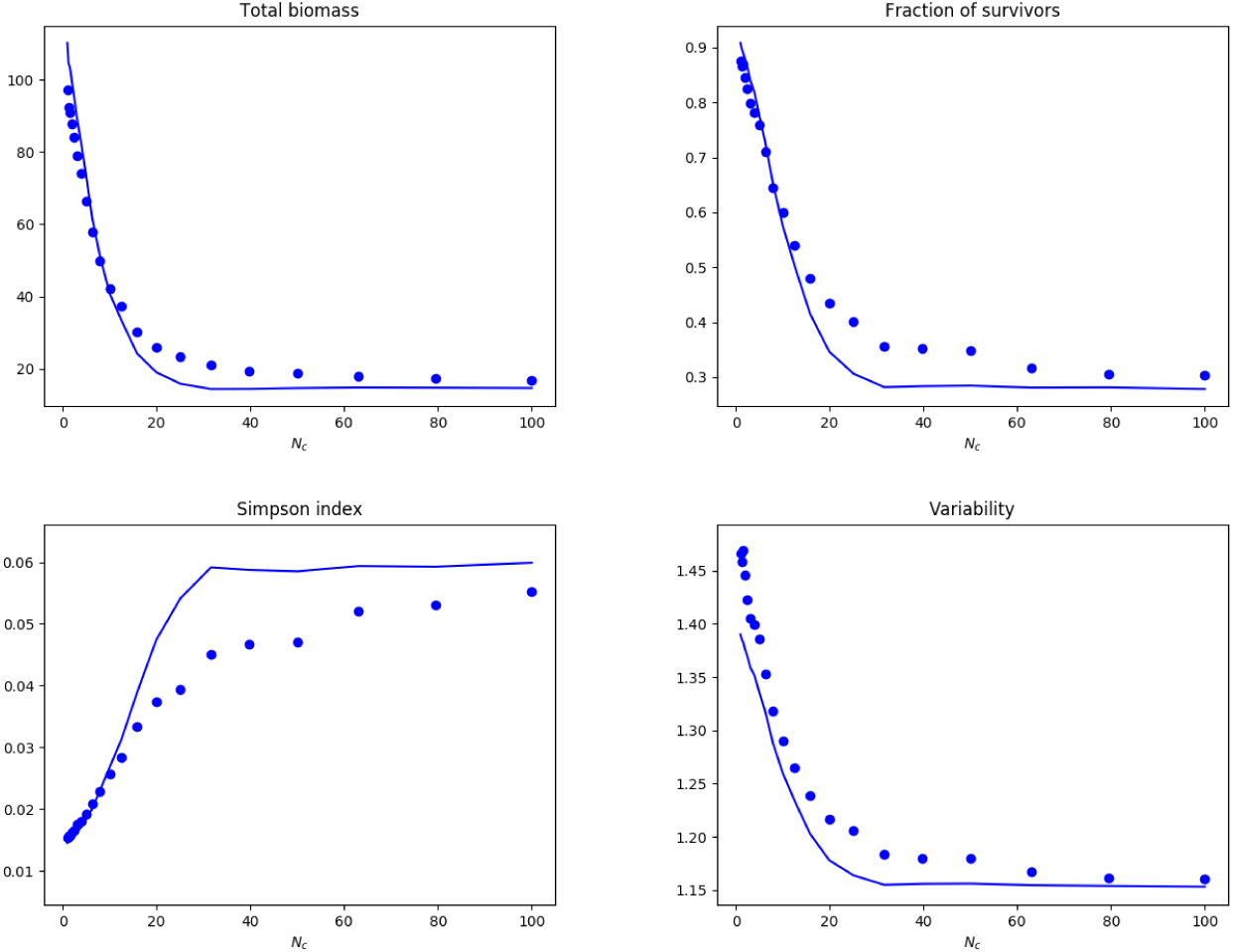
Effect of saturating functional response. Symbols are simulations with a Holling Type II functional response, while solid lines are analytical predictions using the approximate linear-saturating functional response (107).

### 7.2 Heterogeneous means

Let us come back to the cavity method:

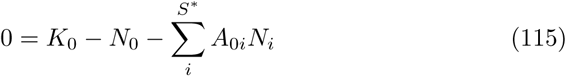

Now let us assume that

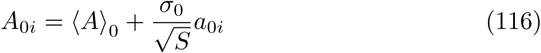

where the mean and the variance both depend on species 0.

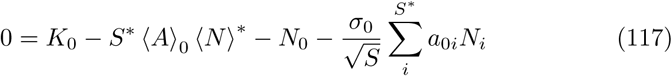

where we still assume that 〈*N*〉 over partners of 0 does not differ from the community average. Then,

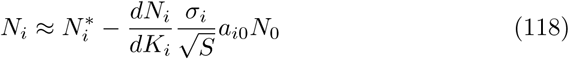

hence

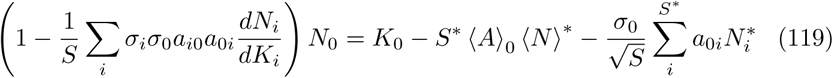

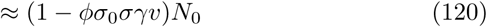

where

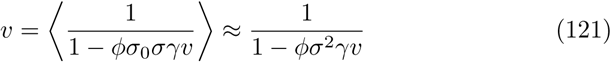

and finally

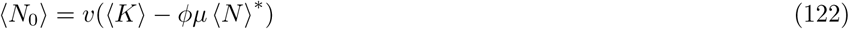

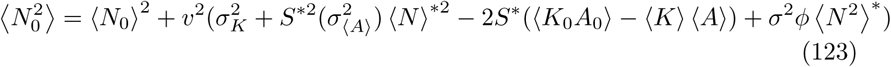

### 7.3 Species groups

The equations can be extended to any structure comprised of discrete groups, with disordered interactions within and between groups, but different statistics for each set of interactions. Coming back to the equilibrium equation, we can write for species *i* in group *x* (which contains *S^x^* species) as

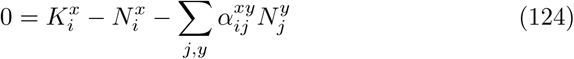

Thus, we now have vector **K** and *ζ* and matrices ***μ, σ, γ,*** defined by

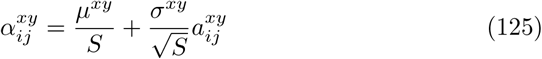

with

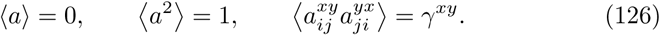

The equations to solve are the same as above, except there are now four equations per group, all coupled: for each group we solve

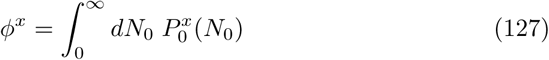

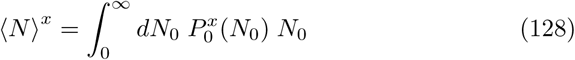

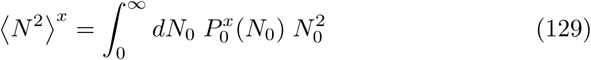

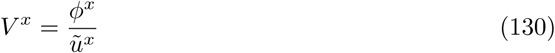

with

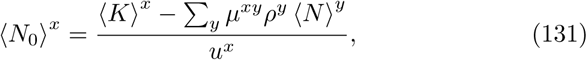

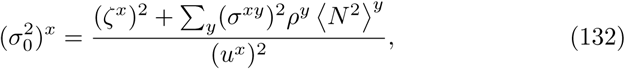

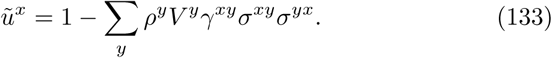

where

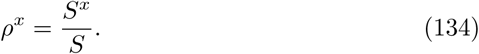

### 7.4 Ordered species

We can take the continuous limit of the mixture model:

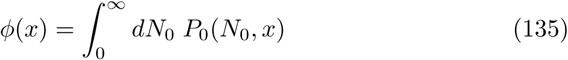

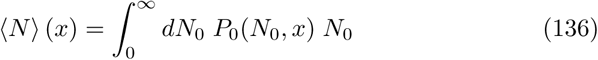

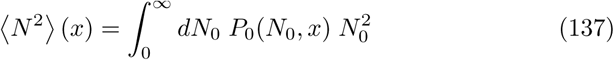

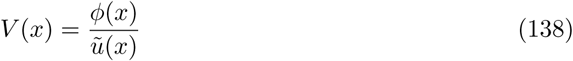

with *P*_0_(*n*, *x*) a Gaussian with mean 〈*N*_0_〉 (*x*) and variance 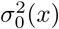, given by

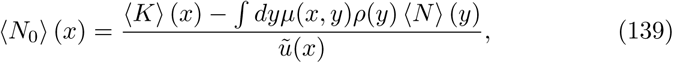

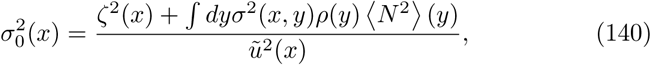

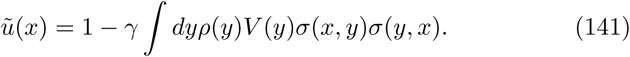

where *ρ*(*x*) is the density of species with rank *x.*

There are two cases that allow for a simple solution: first, if all functions depend only on *x.* Second, if all functions of *x* and *y* depend only on the difference *x − y*, and these functions are either exponential or linear, see results in Fig. 10. By construction, in all these situation *γ*(*x*, *y*) = *γ*(*y*, *x*) must be a constant.

**Figure 10:**
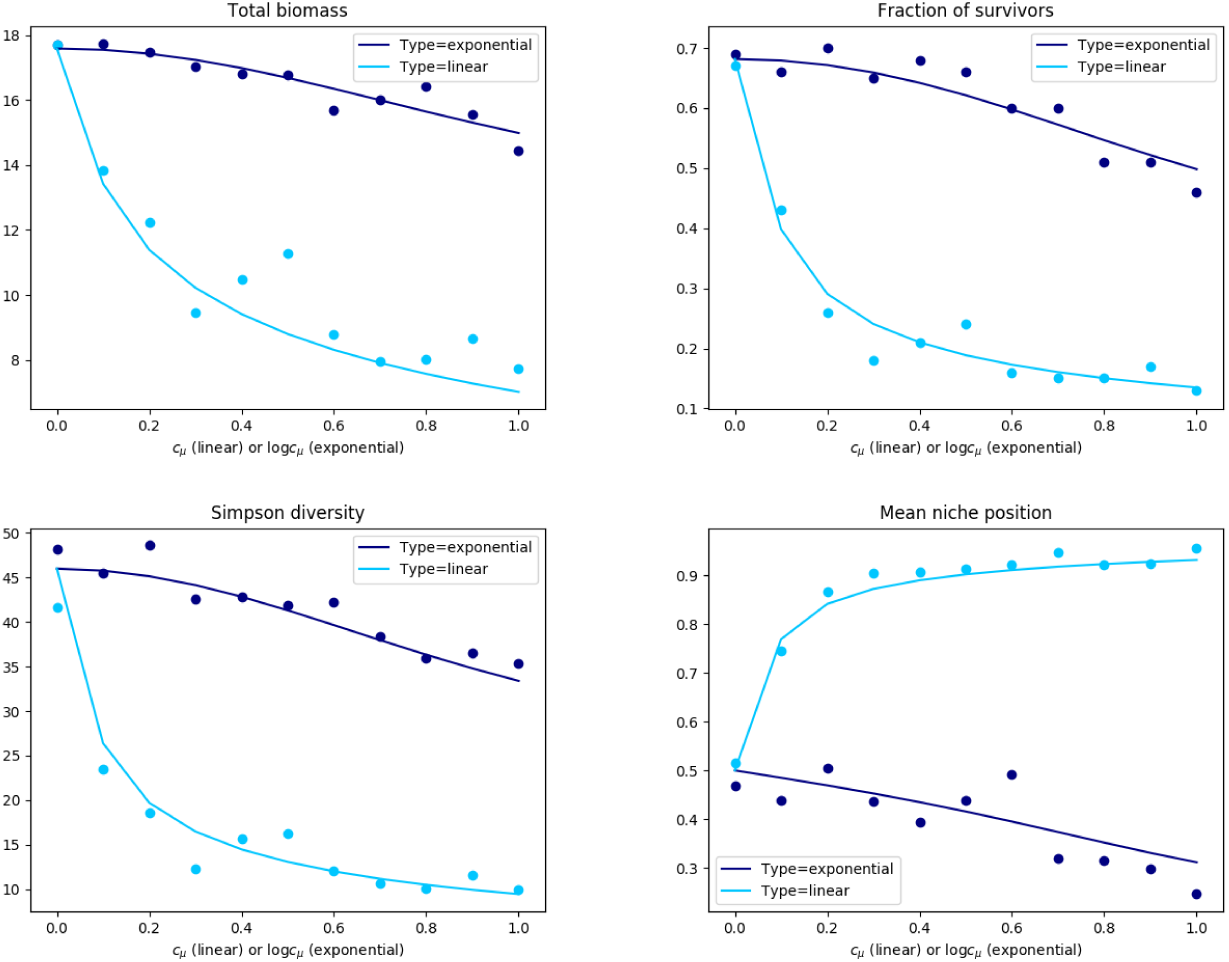
Results for ordered species with exponential and linear scaling. The bottom-right panel is 〈*x*〉*\** the average value of the niche value for surviving species, given that niche values are initially distributed uniformly over [0, 1].

### 7.5 One-sided order

We call one-sided order the situation where functions of *x* and *y* depend only on *x.*

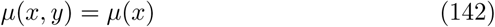

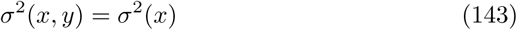

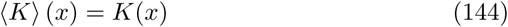

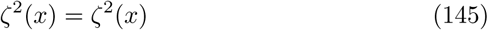

Then, we have

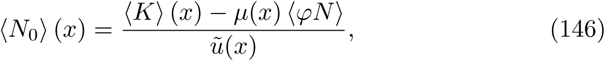

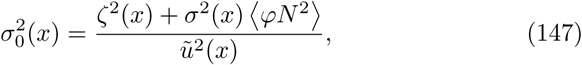

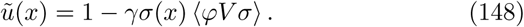

and we can compute

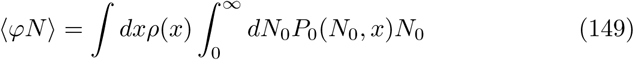

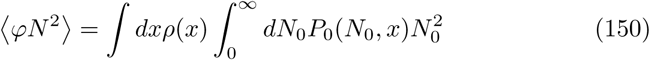

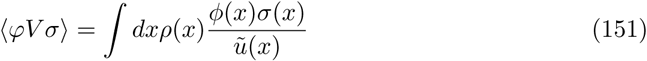

For instance, we can build a tradeoff reminiscent of competition-colonization models by having

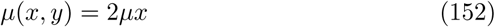

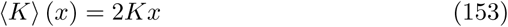

with *μ >* 0 and *σ* and *ζ* constant. In this case, low rank *x* corresponds to high competitive ability (no competition experienced) but low colonization ability.

#### 7.5.1 Exponential order

We can choose exponential functions

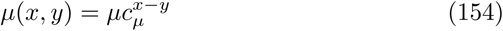

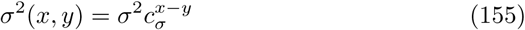

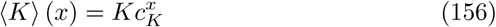

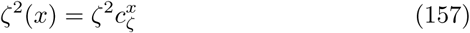

Then, we have

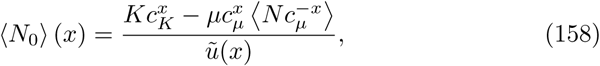

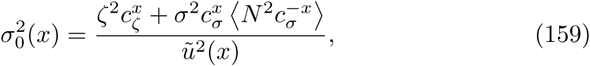

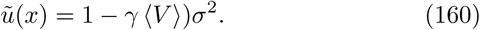

and therefore we must compute

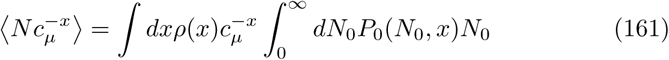

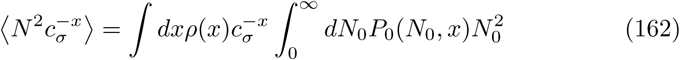

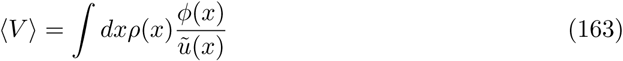

to close the equations. A typical example is metabolic scaling in trophic interactions: given metabolic rate *m_i_* and its scaling with to bodymass *M_i_*, 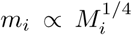, we expect self-interaction *D_i_* ∝ *m_i_* and interaction *A_ji_* ∝ *m_j_*, hence *α_ji_* ∝ *m_j_*/*m_i_* = (1/4)*^x^*^−*y*^ with the niche values given by the logarithm of body mass *x* = log *M_i_* and *y* = log *M_j_*.

#### 7.5.2 Linear order

If we choose linear functions instead

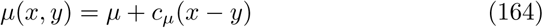

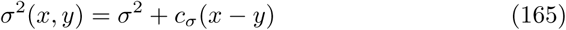

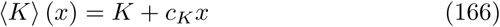

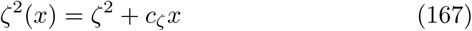

Then we have

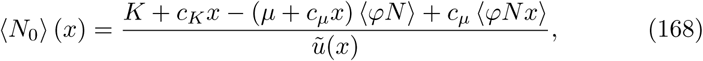

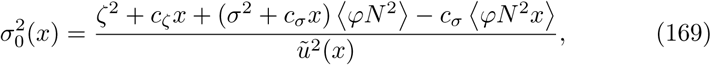

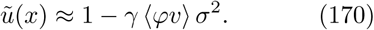

(in fact 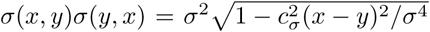 but this makes separation impossible, so we must assume that *c_σ_* is small enough). Hence there are two sets of coupled unknowns: for Ψ = *N*, *N*^2^, *V* we must compute

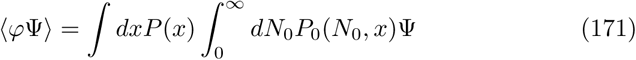

and (except for *V*),

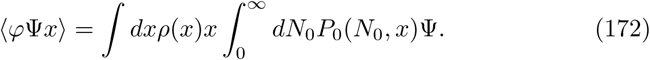

## 8 Appendices

### A. The role of disorder

Computing the equilibrium state in effect requires inverting the matrix *a_ij_* in the equation:

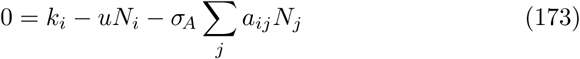

In principle we could expand this equation, substituting

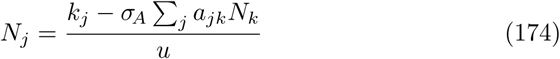

into the sum, and so on recursively, causing all paths *a_ij_a_jk_*… (*i* → *j* → *k…*) in the community matrix to appear in the first equation above. The crucial idea of disordered systems methods is that, for any reasonably complex system, these paths soon lose coherence and start to interfere with each other. These interferences are so complex as to produce seemingly-random results.

One way to see it is that each term in the sum is a (possibly very long) product of coefficients *a_kl_* and is weakly correlated with other such terms – unless the same coefficients appear again and again in the same combinations in many terms. That can happen in a very structured community, or one with only small loops (the extreme is a collection of independent few-species motifs: in each motif, there are not enough paths for them to interfere and create a seemingly random result). This enables the cavity trick, which retains only the first-order feedback of a species on itself, and for the rest needs only the mean and variance of interactions, as the effect of interactions is essentially random.

Let us see a precise example of these ideas. In the main text, we noted that the feedback of a species on itself through the rest of the network is a self-averaging quantity. Consider the following sum:

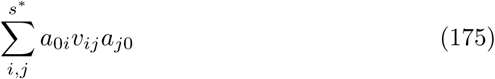

which occurs in (50) (we recall s* = *øs* is the average number of surviving interaction partners for species 0). We separate the sum into two parts:

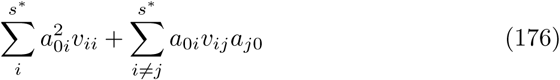

Recall that 〈*a*_0*i*_*a*_*i*0_〉 = *γ* while 〈*a*_0*i*_〉 = 0. Hence, the second sum is a random walk with *s*\*(*s*\* − 1) terms, and its total scales like *s*\*. Now, we should note that

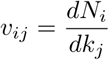

for *i* ≠ *j* is created by the interaction between species *i* and *j,* which is at best proportional to 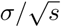 (if the species are linked directly) or even weaker (if they interact only indirectly).

Finally, we get the respective scalings:

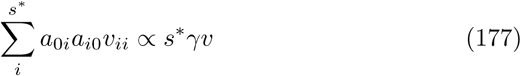

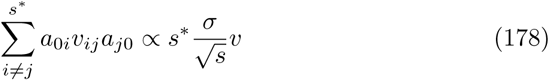

In other words, even though the sum over *i* ≠ *j* has many more terms, it ends up being of order 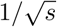 compared to the first sum. That sum itself has variations of order 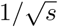 compared to its mean, depending on which species is isolated as species 0. As *s* becomes larger, the feedback thus tends to become a single number, no matter which species we consider.

### B. Linear response

#### Typical cavity method (Press)

Notation: 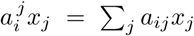. 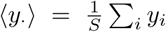. Let *ξ* be a random press-perturbation. Its long term effect on the equilibrium associated to the Jacobian *B* is

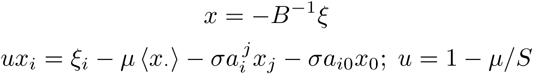

Where we isolated the effect of a generic species 0 (this is the starting point of the cavity method). Assuming a small effect of that species we may write

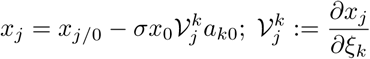

Under the replica symmetry, species 0 is generic and its addition or removal does not change any statistical properties of community, in particular 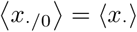. Then

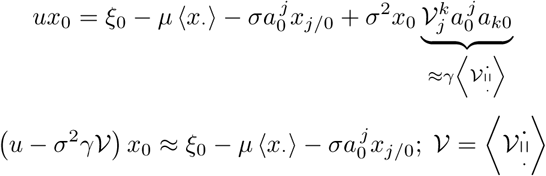

Taking the square

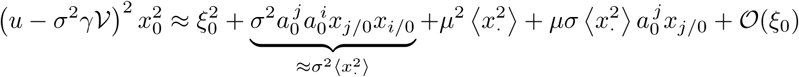

taking the mean over *ξ*_0_, knowing that 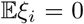;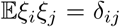, and by linearity, 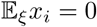, gives

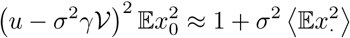

Taking the mean over all species gives

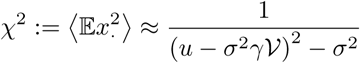

To conclude we need 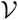. For that we write

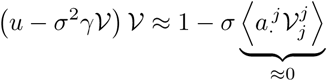

Hence it is solution to (knowing that at *σ* = 0, 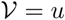)

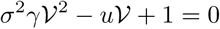

we thus have

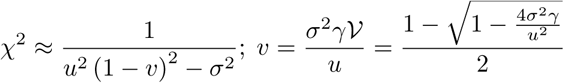

In the Lotka-Volterra assembly perspective, *B* is the interaction matrix and only a fraction *ϕ* of species survive. We still have *u* = 1 − *μ*/*S* but

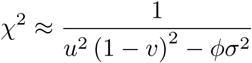

#### Externally applied press

In the assembly perspective, the Jacobian matrix reads

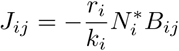

where *r_i_* is the growth rate of species *i* and *k_i_* its carrying capacity. A press acting proportionally to 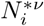 will induce a displacement

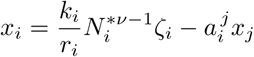

so we can simply adapt the above calculations by replacing *ξ_i_* by 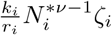. This gives us

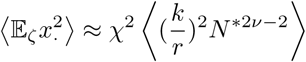

#### To compute (mean case) variability we must solve the Lyapunov equa-tion

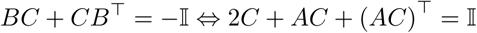

Off diagonal terms, if 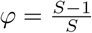, read

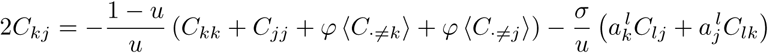

taking the mean, the self-averaging of the terms proportional to *σ* makes the problem reduce to the mean-field case, which gives us the relationship

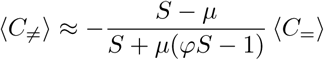

The interaction diversity (*σ*) enters on the diagonal (variances):

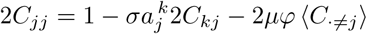

in replacing *C_kj_* in 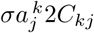 by its expression written above, we only need to track terms proportional to *σ*^2^, the other will self average. So:

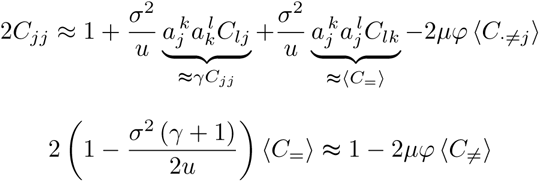

so that

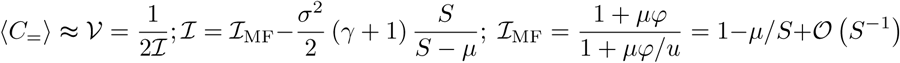

when *μ* = 0, *u* = 1, 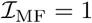. When *σ* = 0, the expression for the mean variance is exact (mean-field case).

#### Demographic noise in the assembly perspective for identical carrying capacities and growth rates

*B* must be replaced with the Jacobian matrix

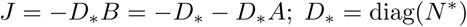

Where the 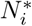 are the equilibrium abundances of surviving species. The Lya-punov equation becomes

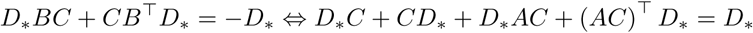

Off diagonal terms if 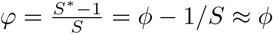, read

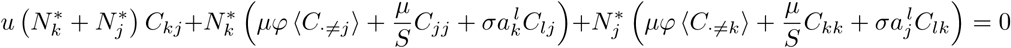

let us write

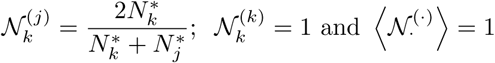

then

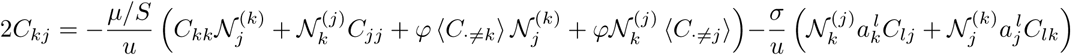

taking the mean over *k* ≠ *j* gives the same approximate relationship between 〈*C*_≠_〉 and 〈*C*_=_〉 as for the mean field case (and is exact when *σ* = 0). On the diagonal:

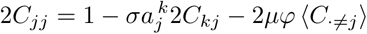

The abundances only enter via the expression of *C_kj_* but always as a ratio 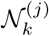 or 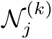 whose average is 1. We get

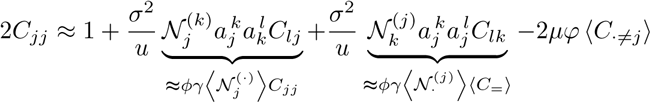

taking the mean over *j* gives

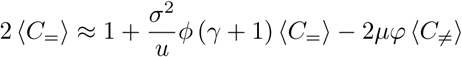

so that

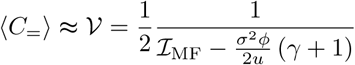

when *μ* = 0, *u* = 1, 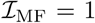. All in all the approximation for (adimensional) invariability is

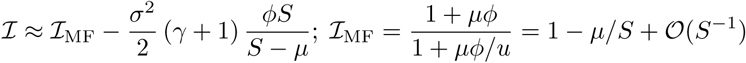

#### More general case in the assembly perspective

In general *B* must be replaced with the Jacobian matrix

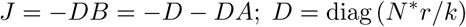

Under stochastic perturbations scaling as 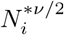 (*v* = 1 demographic noise, *v* = 2 environmental noise) the Lyapunov equation becomes

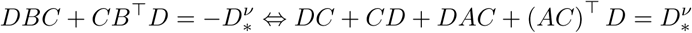

For off-diagonal terms we may reproduce the previous calculations by replacing 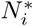 by 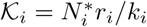 and

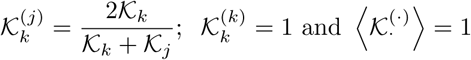

On the diagonal however we now have:

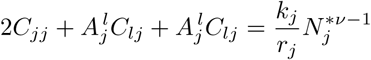

which, as above, leads us to

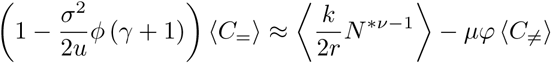

so that

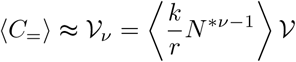

In particular, for *v* = 0, 1, 2 (immigration, demographic and environmental noise, respectively):

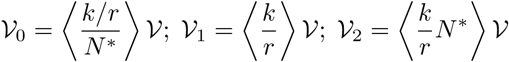

#### Taylor’s law

relates species mean abundance and their variance. In our context, under some stochastic noise acting independently on species (mean case scenario) we have that

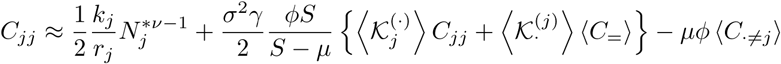

We roughly approximate 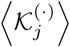 and 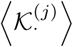 by 1. Using that

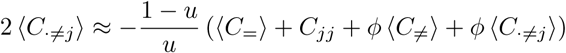

so

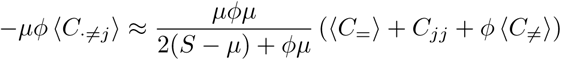

If we define

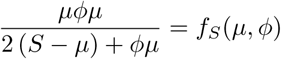

This leads us to

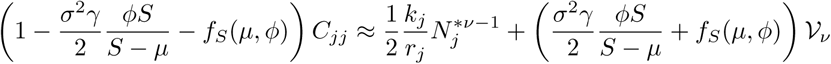

if *μ* is of order zero in 1/*S* then *f_S_* is of order one and for *S* large enough

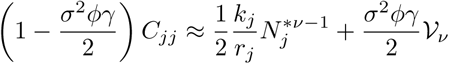

We thus expect an exact power law 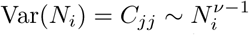 when *γ* = 0 and *σ^2^* large enough (so that *N_j_* and *k_j_* become uncorrelated).

### C. Other calculations

#### Generating *A* with correlation to *K*

Here is how to generate *A_ij_* with prescribed correlation to *K_i_*:

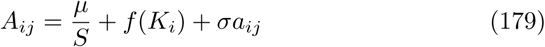

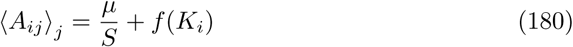

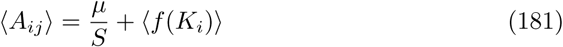

Let’s set 〈*K_i_*〉 = 1, then:

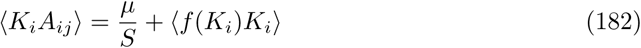

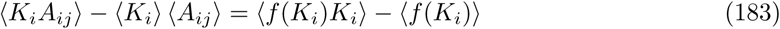

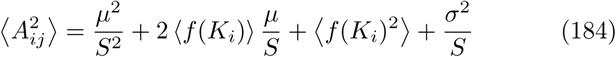

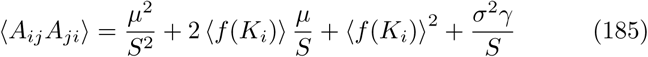

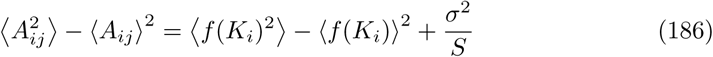

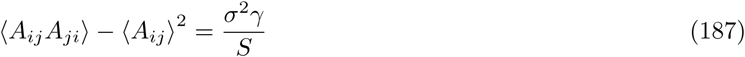

Now, let’s set *f*(*K_i_*) = *c*(*K_i_* − 1) so that

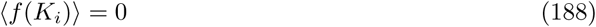

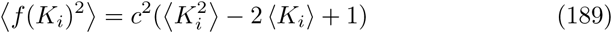

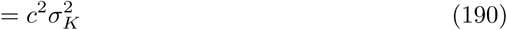

Then

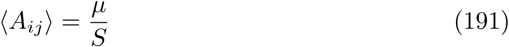

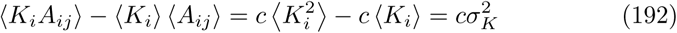

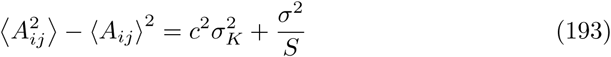

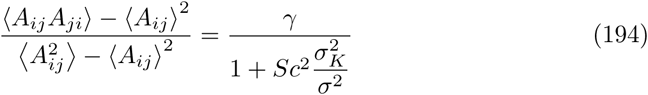

Hence, achieving a prescribed correlation 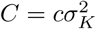 requires to generate a matrix *a_ij_* with parameters

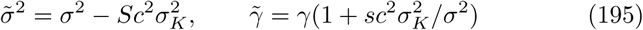

before inserting it into 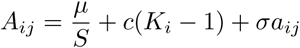.

## Footnotes

1 It became prominent with the study of disordered materials such as glasses [29], before being applied to algorithmics and information theory [28], chemistry [30], economics [33] and systems biology [35].

2 The term of “equilibrium” is ambiguous as it has been used for the Theory Island Biogeography among others. A fully stochastic system can indeed be characterized by probability distributions that attain a stationary state – in physics, this is called a Non-Equilibrium Stationary State, or driven stationary state, to distinguish it from the concept of equilibrium in deterministic dynamics and from thermodynamic equilibrium.

3 Even with many local attractors, this picture may remain valid if the dynamics exhibit *aging,* i.e. the sytem is prone to falling into deeper and deeper attractors as time passes.

4 Note however that in Sec. 7.1, we extend these results to a saturating functional response.

5 The reader may recognize this idea as similar to that of moment equations, see e.g. [8].

6 Note that keeping the same *s* for different pool sizes *S* means that connectivity *c* = *s/S* must decrease with pool size.

7 It can be achieved with a power-law tail whose exponent is around or below 2, meaning that the variance of the theoretical distribution is arbitrarily large or even diverges, but in effect, a few massive outliers will be responsible for this variance, while most interactions will be negligible.

8 In fact, this is a useful parametrization, since the Lotka-Volterra approximation to more realistic dynamics may not hold if couplings are too large *|αij| >* 1 (e.g. mutualistic inter-actions would then cause abundances to diverge). Hence couplings also give an idea of the range of validity of the model.

9 A classic case is Levins and Culvers’s colonization-competition tradeoff [19] where only a very special and asymmetrical structure ensures the survival of more than a single species. We discuss it further in [4].

10 This assumption is correct if there is only one globally stable assembled state; the method should later be extended to settings with alternative stable states, where the addition of a new species could cause a sudden shift between different equilibria.

11 Further clarifications on how randomness intervenes in interactions can be found in Appendix A.

12 This assumption was found to be essential to allow high coexistence.

